# In situ cryo-ET reveals restricted docking of intraflagellar transport at the base of the trypanosome flagellum

**DOI:** 10.64898/2025.12.23.690766

**Authors:** Sophia R. Staggers, Mingqi Zhao, Oliver Harrigan, Sabrina Absalon, Yameng Huang, Zhangyu (Sharey) Cheng, Yongxin (Leon) Zhao, Muyuan Chen, Stella Y. Sun

## Abstract

Cilia and flagella are conserved organelles essential for sensory and motility functions and are assembled and maintained by bidirectional intraflagellar transport (IFT) along the axoneme. In *Trypanosoma brucei*, IFT is concentrated at the flagellar base near the transition zone and is restricted to four microtubule doublets (MTDs) as it traverses from the transition zone to the axoneme before reaching the extra-axonemal structure, the paraflagellar rod (PFR), associated region. To gain an insight on its initial working model before IFT restriction occurs, we revealed the in situ architecture of the IFT complex during the initial and complete landing to the transition zone using cryo-focused ion beam (cryo-FIB) milling and cryo-electron tomography (cryo-ET). IFT proteins assemble into polymeric trains upon landing. However, in the proximal portion of the flagellum, subtomogram averaging revealed that each single train is associated with individual MTDs at variable A- or B-tubule positions. By using miniaturized and enucleated *T. brucei* zoids that preserve the full circular arrangement of MTDs without milling, we revealed an alternating but spatially restricted IFT train pattern during the landing process, resembling the anterograde train scaffold known in *Chlamydomonas*. Magnify expansion microscopy further confirmed this restricted distribution of IFT proteins at the basal pool. Depletion of the IFT component IFT46 protein reduced the number of simultaneously docked trains without altering the restricted landing spatial pattern. These findings reveal a previously unrecognized spatial regulation governing IFT train initiation at the flagellar base, which could operate independently of IFT-B structural integrity, bridging the gap between the IFT architecture and doublet-spatial docking prior to anterograde transport.

## Introduction

Cilia or flagella are evolutionarily conserved cellular structures that perform crucial sensory and motility functions, and their assembly is powered by intraflagellar transport (IFT). IFT was first identified in the green alga *Chlamydomonas reinhardtii* and is now characterized as a conserved bidirectional transport system that shuttles protein complexes and cargoes along the axonemal microtubules doublets (MTDs) (Kozminski et al., 1993, Kozminski et al., 1995). Anterograde IFT, from the cell body to the flagellum tip, is driven by kinesin-2 motors, while retrograde IFT, from tip to base, is powered by dynein-2 motor proteins (Kozminski et al., 1995, Pazour et al., 1998, Porter et al., 1999, Kozminski et al., 1993). IFT train cargoes and motor proteins are assembled upon a scaffold composed of two repeating multiprotein complexes, with the 16-subunit IFT-B complex forming the core of anterograde trains and the 6-subunit IFT-A complex forming the core of retrograde trains in *Chlamydomonas* (Jordan and Pigino, 2021, Cole et al., 1998, Piperno and Mead, 1997). At the base, a specialized connecting structure known as the transition zone has long been believed to serve as a structural and regulatory gate that controls the entry or exit of proteins, including the IFT complexes to maintain ciliary or flagellar composition (Gilula and Satir, 1972, Breslow et al., 2013, Lin et al., 2013, van den Hoek et al., 2022, Kee et al., 2012). Recent work in *Caenorhabditis elegans* has further demonstrated that the transition zone is not just a passive structural region, but also a physical barrier that IFT complexes must be actively driven through by dynein motors (De-Castro et al., 2022). Disruption of IFT components or their regulators causes a diverse group of diseases known as ciliopathies, which can be devastating to proper function of the brain, heart, skeleton, and other organs (Reiter and Leroux, 2017).

To understand how this conserved machinery adapts to specialized flagellar architectures, *Trypanosoma brucei* has emerged as a powerful model for studying IFT asymmetry and doublet selectivity (Bertiaux et al., 2018a, Araujo Alves et al., 2025). *T. brucei* is responsible for African trypanosomiasis in humans and Nagana in animals and relies on a single motile flagellum that plays essential roles in cell motility, morphogenesis, and host transmission via the tsetse fly vector. The flagellum emerges through the flagellar pocket (FP), which encloses the transition zone, and is mechanically anchored to the cell body through the flagellar attachment zone (FAZ) (Kohl and Gull, 1998, Sherwin and Gull, 1989). This coupling enables the twisting and constrictive movements characteristic of *T. brucei* motility, facilitating passage through confined environments (Sun et al., 2018, Rodríguez et al., 2009). Despite its extensive morphological remodeling throughout its complex life cycle, *T. brucei* retains a conserved 9+2 axonemal architecture, with each MTD consisting of a complete A-tubule and an incomplete B-tubule arranged in a B–A orientation when viewed from the proximal flagellar base (Ralston et al., 2009, Langousis and Hill, 2014). Beyond its conserved axonemal structures, the *T. brucei* flagellum features an extra-axonemal structure, the paraflagellar rod (PFR), which runs parallel to the axoneme and affixes to MTDs 4-7 through an intricate network of connecting proteins essential for motility (Vickerman, 1962, Farina et al., 1986, Anderson and Ellis, 1965, Cachon et al., 1988, Bastin et al., 1998, Lacomble et al., 2009). Such structural asymmetry is similar to that of mammalian sperm flagella and primary cilia, where specialized doublet architecture is critical for motility and sensory functions, and defects lead to infertility and ciliopathies (Inaba and Mizuno, 2016, Pariz et al., 2021). The asymmetric positioning of the PFR highlights the asymmetry in *T. brucei* flagellum construction, which is further underscored by the distinctive and highly asymmetric distribution of IFT complexes in *T. brucei*. Focused-ion beam-scanning electron microscopy (FIB-SEM) studies of IFT in the insect form of *T. brucei* revealed that IFT trafficking is restricted to two discrete pairs of MTDs in the axoneme, only appearing near doublets 3-4 and 7-8, positioned opposite the PFR (Bertiaux et al., 2018a). Yet the basis for this doublet selectivity remains unresolved, including whether anterograde and retrograde trains traffic exclusively on a distinct A- or B-tubule of MTDs, and it raises a fundamental question: how is IFT first organized at the flagellar base, particularly at the transition zone, where trains can be assembled and docked before entering the axoneme.

At the flagellar base, IFT proteins and motors are highly concentrated in a large basal pool in *C. reinhardtii*. Cryo-ET and ultrastructure expansion microscopy (U-ExM) revealed that anterograde IFT trains assemble and dock on all nine MTDs at the transition zone, initiating with polymerization of the IFT-B scaffold followed by recruitment of IFT-A and dynein complexes (van den Hoek et al., 2022). In *T. brucei*, however, the flagellum is stable, where mature flagella do not undergo continuous turnover like those in *Chlamydomonas*, suggesting fundamentally different IFT regulatory strategies. Recent high-resolution fluorescence and FIB-SEM imaging demonstrated that this selectivity occurs at the proximal flagellum, as IFT trains emerge from the FP, where they switch among multiple microtubules, a behavior also recently discovered in primary cilia (Araujo Alves et al., 2025, Sun et al., 2025). However, the molecular and structural mechanisms that organize IFT at the basal pool, and whether trains are assembled during their transition through the transition zone remain unknown. In particular, it is unknown whether *T. brucei* establishes an initial landing selectivity at the transition zone that precedes the well-characterized doublet restriction in the axoneme.

Our study focused on mature flagella in non-dividing *T. brucei* cells. Two potential models of IFT train assembly and doublet selection were considered: First, assembly at the axoneme–PFR interface followed by redistribution onto selective tracks; Second, assembly immediately before flagellar entry followed by switching to restricted doublets. To distinguish between these possibilities, we examined the transition zone, recently identified in *Chlamydomonas* as the primary site of IFT train assembly and docking (van den Hoek et al., 2022). Using cryo-ET on cryo-FIB-milled *T. brucei*, we observed polymeric IFT trains preassembled in the cytoplasm near the transition zone, and undergoing both initial and complete landing while aligned parallel to the transition zone MTDs within the FP. Because capturing these transient events was inconsistent in single-flagellated wild-type cells using cryo-FIB-milling, we employed genetically engineered enucleated “zoids,” derived from Centrin4 knockdown mutants (Shi et al., 2008, Sun et al., 2018). The use of zoids enabled reproducible visualization of IFT train initial landing at the flagellar base in situ. Machine-learning based segmentation showed that anterograde trains did not simultaneously occupy all nine MTDs within the transition zone. Instead, only a limited number of trains were present at transition zone per cell, each associated with either the A- or B-tubule and distributed asymmetrically to opposite sides of the transition zone following complete landing. Subtomogram averaging of IFT densities revealed an anterograde-type scaffold architecture near the transition zone consistent across cells. Depletion of IFT46, a structural component of the IFT-B complex, reduced the number of simultaneously docked trains but did not alter their asymmetric spatial distribution or the absence of a preferred landing position along the trafficking axis. These datasets provide the first in situ model of IFT train landing in *T. brucei*. Understanding this process is crucial for determining whether an initial selectivity step exists before IFT selectivity occurs along the axoneme, potentially revealing a unique, trypanosome-specific mechanism of IFT initiation at the basal pool.

## Results

### Characterization of IFT train initiation at the flagellar base

Flagella are microtubule-based organelles composed of an axoneme extending from the basal body, where IFT proteins concentrate. To visualize if IFT proteins initiate train assembly at the transition zone of the flagellum base, an area deeply embedded within the cell body, we performed cryo-FIB-milling to thin wild-type *T. brucei* cells and generate thin lamellae suitable for cryo-ET (Fig. 1A and S1A). Tilt series were collected from these lamellae using cryo-electron microscopy and reconstructed into tomograms, revealing the three-dimensional architecture of the flagellum base. A total of 66 lamella, ranging in thickness from 160 to 250 nm, were prepared. Among these, 54 tilt series were successfully acquired and reconstructed into tomograms that included the flagellum base, out of which 21 tomograms contained visible IFT train-like densities (Fig. S1B). Only 3 tomograms preserved all nine MTDs due to the challenge of precise depth control while milling (Fig. S2). 2D tomogram slices revealed distinct IFT train–like densities at the base of the flagellum, within the asymmetric membrane-invaginated FP (Fig. 1B). 3D feature annotation was performed through the full tomographic volume, identifying six major structural components: the basal body, transition zone, basal plate, axoneme, FP membrane and IFT trains (Fig. 1B’ and S2 and Movie S1).

**Figure 1:**
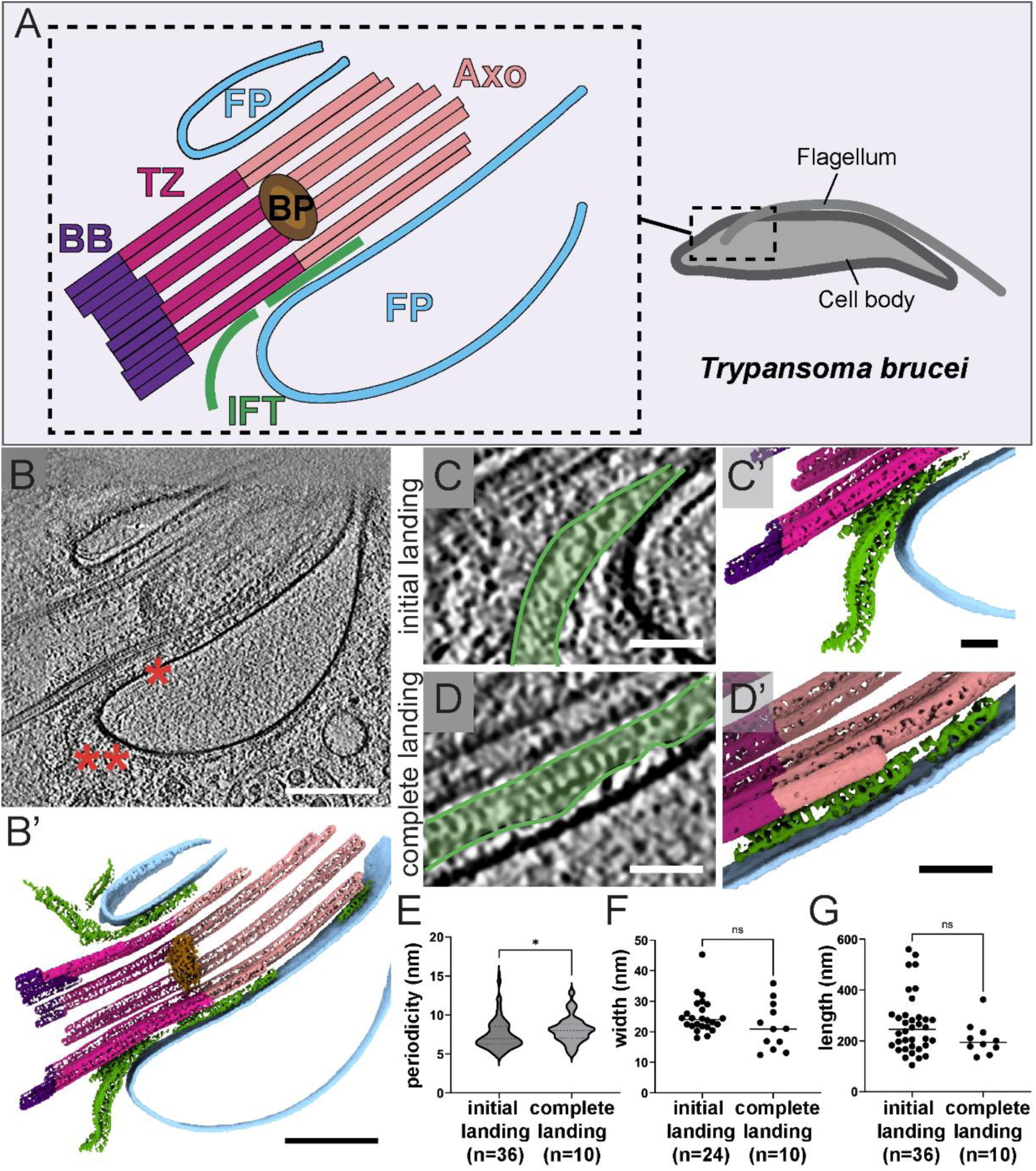
Identification of two landing stages of IFT trains in wild-type *T. brucei*. **(A)** Schematic of a *T. brucei* cell highlighting the flagellar base region (dashed box), featuring key structures involved in IFT including flagellar pocket (FP), axoneme (Axo), transition zone (TZ), basal body (BB) and basal plate (BP). **(B)** 2D tomogram slice of the flagellum base, indicated by the curved membrane of the flagellar pocket (FP), featuring densities of IFT trains at two different landing stages including initial landing (single red asterisk), in which the front end of the train is touching the microtubule doublet (MTD), and complete landing (double red asterisks), in which the whole train is parallel along the MTD. Scale bar 200 nm. **(B’)** 3D annotation of the tomogram in (B) showing the axoneme (light pink), transition zone (magenta), extended from the basal body (purple) at its proximal end and ended at the plane of the basal plate (brown) at its distal end, FP membrane (light blue), and IFT (green). **(C-D)** Zoomed-in tomogram slices highlighting (C) initial and (D) complete landing trains (red asterisks as in A). **(C’-D’)** Zoomed-in corresponding 3D annotations of B and C. Scale bar 50 nm. **(E)** Periodicity measurements of initial (7.48 ± 1.93, n = 36) and complete (8.12 ± 1.69, n = 10) landing trains. **(F)** Width of initial (25.3 ± 5.83 nm, n = 24) and complete (21.8 ± 7.67 nm, n = 10) landing trains measured from 2D slices were similar. **(G)** Length of initial (265 ± 118 nm, n = 36) and complete (208 ± 65.5 nm, n = 10) landing trains were similar.

The FP membrane partially aligns with the transition zone, which allows us to distinguish the train landing process at different stages. Based on the spatial arrangement of IFT-like train densities, we define two sequential stages of train association at the flagellum base. The initial landing stage is characterized by a curved IFT train whose front end is touching the transition zone (Fig. 1C, C’). This configuration resembles the early assembly stage of IFT trains in *C. reinhardtii* observed by cryo-ET and subtomogram averaging, where immature trains display a characteristic curved “tail-like” morphology (van den Hoek et al., 2022). To quantify train positioning, we measured the distance from the contact point of each initial landing train to the basal body and normalized it to the total transition zone length. The resulting ratio ranged from 0.13 to 0.77, indicating that initial IFT train landings occurred throughout most of the transition zone without a single preferred site (Fig. S8A and B). Next, the complete landing stage is defined by trains fully aligned and parallel to the MTDs of the transition zone and/or axoneme (Fig. 1D, D’).

Structural and imaging studies can reveal distinct architectures between anterograde and retrograde IFT trains. Cryo-ET studies in *C. reinhardtii* demonstrated that IFT trains undergo a major structural rearrangement during the transition from anterograde and retrograde transport. In the anterograde transport, trains display a periodic spacing of ~6.5 nm between IFT-B subunits and ~11.5 nm between IFT-A subunits, whereas retrograde trains adopt a ~45 nm repeat after structural rearrangement at the flagellar tip (Jordan et al., 2018, Lacey et al., 2023, Lacey et al., 2024). In *T. brucei*, measurements from our tomograms revealed a mean periodicity of 7.48 ± 1.93 nm during the initial landing stage and 8.12 ± 1.69 nm during complete landing (Fig. 1E). These values closely match the anterograde IFT patterns and differ from the 45 nm spacing characteristic of retrograde trains (Jordan et al., 2018, Lacey et al., 2024), confirming that these observed train densities correspond to anterograde IFT, revealing that their entry occurs stochastically along the transition zone rather than fixed docking sites.

Previous live-cell fluorescence microscopy reported average IFT train lengths of 393 ± 51 nm for anterograde and 250 ± 51 nm for retrograde transport in *T. brucei*, suggesting that the IFT trains undergo a length-regulation step prior to the anterograde switch to retrograde at the flagellar tip (Buisson et al., 2013). It remains unclear whether a similar regulatory step occurs earlier, specifically during the transition from cytoplasmic assembly to flagellar entry after complete docking at the base. To assess potential changes in train architecture at this entry point, we quantified the length and width of IFT trains in both the initial landing and complete landing stages. The average train width remained relatively consistent with 25.3 ± 5.8 nm during initial landing and 21.8 ± 7.7 nm during complete landing (Fig. 1F). In contrast, the train length was characterized from 265 ± 118 nm in the initial landing to 208 ± 65.5 nm in the complete landing trains on average, showing slightly shorter, but insignificantly different length with less variation by cryo-ET (Fig. 1G). In cryo-FIB–milled lamellae, we consistently observed a maximum of four anterograde IFT trains assembled and docked at the transition zone (Fig. 4C). Even in tomograms where all nine MTDs were preserved, IFT trains were never distributed across every doublet (Fig. S2). Within our dataset, the largest number of simultaneously docked trains at the proximal transition zone was three. Together, these observations indicate that *T. brucei* assembles and docks only a limited number of anterograde IFT trains at the flagellar base, suggesting a spatially constrained or sequential initiation process. Further investigation using fully preserved transition zone architectures and reproducible whole-cell cryo-ET will be essential to reveal this finding.

### IFT trains associate with individual MTDs at transition zone-axoneme interface

The nanometer resolution achieved by cryo-ET is sufficient to distinguish adjacent MTDs, a detail that remains challenging by using conventional dual-beam SEM and FIB imaging (Bertiaux et al., 2018a, Alves et al., 2024, Araujo Alves et al., 2025). This resolution of detail allowed us to precisely examine IFT–MTD association, addressing whether IFT trains attach to individual MTDs or bridge adjacent doublets, which is a key question given the restricted IFT localization to paired MTDs on one side of the axoneme. In a representative cryo-FIB-milled tomogram of wild-type cell, 3D segmentation identified a total of three IFT trains at the complete landing stage, two of which were positioned on adjacent MTDs (Fig. 2A). Combined with their characteristic anterograde periodicity, this spatial organization revealed that two anterograde IFT trains can simultaneously associate with neighboring MTDs at the interface between the transition zone and axoneme. This observation is consistent with recent super-resolution fluorescent microscopy discoveries showing that IFT trains associate to individual MTDs (Araujo Alves et al., 2025), while extending these findings by visualizing assembled train structure in situ using cryo-ET.

**Figure 2:**
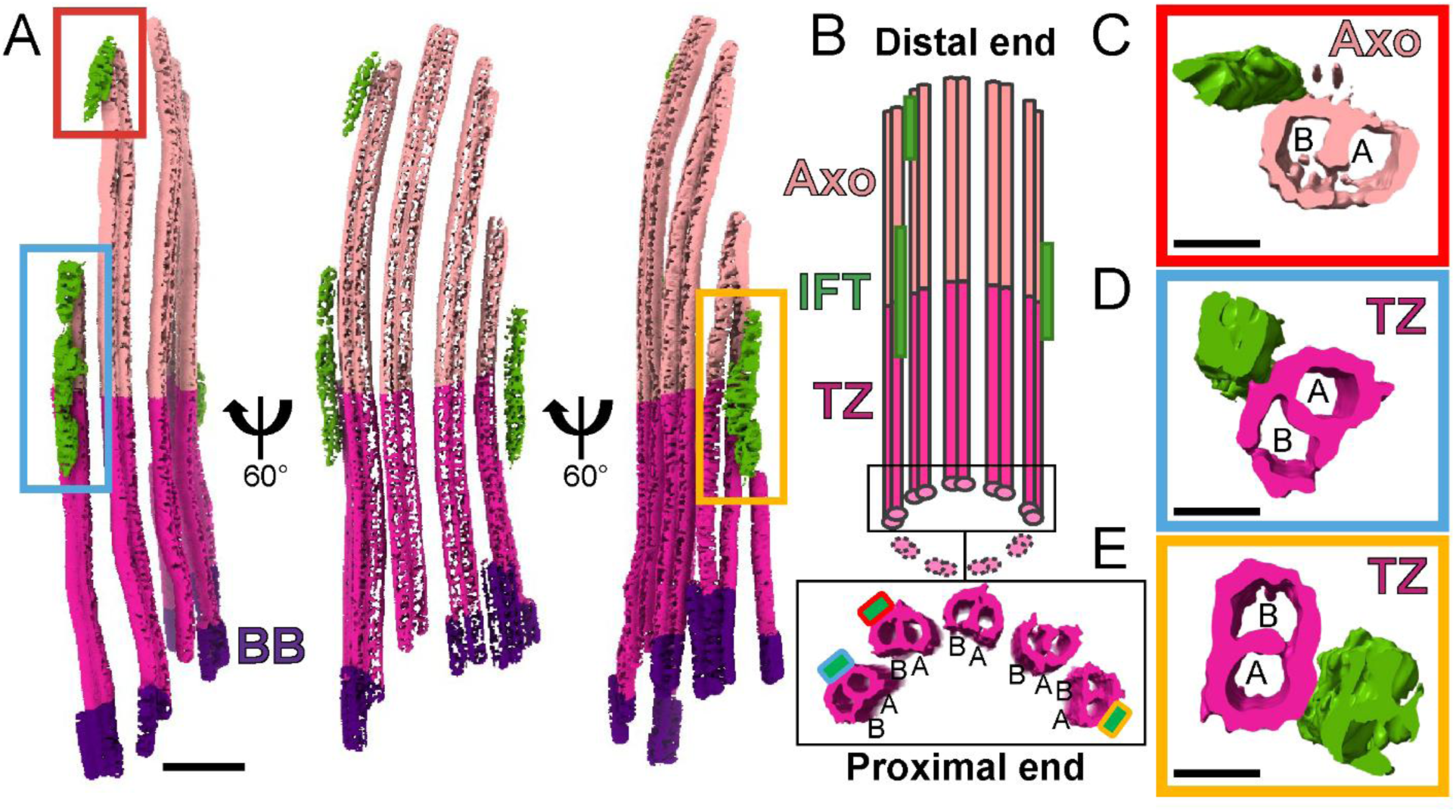
IFT trains associate with both A and B tubules of microtubule doublets. **(A)** 3D annotation of flagellum tomogram from Fig. 1B featuring three complete landing anterograde IFT trains (green), distributed along the interface between transition zone (magenta) and axoneme (pink) before PFR initiates. Scale bar 100 nm. **(B)** Schematic illustration of IFT train distribution along the axoneme (Axo) and transition zone (TZ). Subtomogram averaging of the MTD yielded a map at ~30.5 Å resolution within the transition zone and a map at ~29.5 Å resolution within the axoneme. Mapping back of the averaged segments to the tomogram revealed the configuration of A- and B-tubules from the proximal end zoomed-in in the black box. Small green rectangles within the black box represent IFT train association to MTDs, with outline colors corresponding to boxed trains in (A). Three distinct association regions were identified including **(C)** the trains bound to the B-tubule only, **(D)** trains at the B- and A-tubules interface, and **(E)** trains bound to the A-tubule only. Scale bar 20 nm.

Correlative total internal reflection microscopy and electron tomography within *C. reinhardtii* demonstrated that individual MTDs can accommodate bidirectional transport. Specifically, anterograde trains travel along the B-tubule, while retrograde trains return along the A-tubule (Stepanek and Pigino, 2016). To determine whether these anterograde trains docked on the A- or B-tubule in *T. brucei*, we performed subtomogram averaging of the transition zone MTDs, generating a ~31 Å structure across a ~66 nm segment, which provided sufficient resolution to distinguish the complete A-tubule cylinder from the incomplete B-tubule (Fig. 2D and E). A comparable averaging of axonemal MTDs generated a ~30 Å structure across a ~67 nm segment (Fig. 2C). Mapping these averaged segments back into the tomographic volume allowed us to assign A- or B-tubule identity within the circular MTD arrangement (Fig. 2B). Our analysis revealed one IFT train associated between the B- and A-tubules of a single doublet within the transition zone, a second associated to the B-tubule of an adjacent doublet, and a third docked to the A-tubule on the opposite side (Fig. 2B-E). These findings demonstrate that anterograde trains in *T. brucei* can associate with both A- and B-tubules, in contrast to *C. reinhardtii,* where anterograde trains are confined exclusively to B-tubules (Stepanek and Pigino, 2016). This suggests that in *T. brucei*, tubule identity alone does not dictate the directionality of IFT trafficking, indicating a distinct mechanism of anterograde train engagement at the flagellar base.

### Characterization of IFT distribution around the flagellar base in enucleated zoids

While cryo-FIB milling enabled nanometer-resolution localization of individual IFT trains on individual MTDs within the transition zone, approximately 95% of the tomograms targeting the flagellar base showed partial lateral loss of the transition zone. This loss compromised our ability to analyze the full spatial organization of IFT trains around the transition zone landing site. This highlights the need for a more efficient thinning method to study IFT spatial dynamics at the flagellar base. Enucleated *T. brucei* mutants, known as zoids, previously shown to be thin enough for whole-cell cryo-ET without FIB milling, provided a promising alternative (Sun et al., 2018, Bregy et al., 2023). Zoids retain motility, viability, and flagella length while exhibiting a reduced cellular volume that fits within the optimal imaging range of cryo-ET. These zoids allow direct visualization of the flagellar base and enable quantitative analysis of the frequency and spatial distribution of IFT trains without requiring physical milling.

Before proceeding with cryo-ET on zoids, we needed to evaluate whether the IFT concentration and distribution at the flagellar base differ between zoids and wild-type cells. We endogenously tagged the N-terminus of IFT81 or IFT88 with eGFP using the PCR-only tagging (pPOT) system via Centrin4-RNAi induction (Dean et al., 2015, Paterou et al., 2025). Centrin4 depletion in each cell line was confirmed by reverse transcription PCR (RT-PCR) (Fig. S3A and B), and western blot analysis with anti-GFP antibodies confirmed that IFT81 and IFT88 expression levels remained unchanged upon Centrin4 depletion (Fig. S3C and D). Based on growth assay of Centrin4-RNAi and percentage of zoids in the population at each time point over 96 hours of tetracycline induction, we determined 72 hours as the optimal induction timepoint to isolate zoids (Fig. S4). To assess whether IFT levels differ between wild-type and zoid cells, we performed confocal z-stack imaging and quantified autofluorescence intensities of eGFP::IFT81 and eGFP::IFT88 in Centrin4-RNAi induced zoids and uninduced wild-type controls. Because zoids lack a nucleus and do not divide, our analysis focused on their mature flagella and compared them to mature flagella from non-dividing wild-type cells at the one kinetoplast, one nucleus (1K1N) stage identified by DAPI staining (Fig. S5A). Fluorescence intensity measurements revealed no significant differences in IFT concentration between the two groups (Fig. S5B). For eGFP::IFT81, the average fluorescence was 131.8 ± 19.2 A.F.U. in wild-type cells (n = 40) and 141.5 ± 28.6 A.F.U. in zoids (n = 40). Similarly, for eGFP::IFT88, the averages were 162.4 ± 36.3 A.F.U. in wild-type cells (n = 40) and 150.7 ± 30.4 A.F.U. in zoids (n = 40). These results indicate that IFT concentration at the base of the mature flagellum is comparable between wild-type cells and zoids.

Because conventional confocal microscopy cannot reliably distinguish one versus two closely spaced trains when small landing gaps are present, we next compared the spatial distribution of IFT proteins at the mature flagellar base in wild-type cells and zoids using the recently developed Magnify Expansion Microscopy (ExM) (Klimas et al., 2023). This method incorporates simultaneous biomolecular anchoring and gelation with a reinforced hydrogel, achieving up to ~11x expansion for enhanced protein localization. The measurements of axoneme diameters confirmed average expansion factors of 7.2x in wild-type cells and 6.7x in zoids (Fig. S6D). Immunofluorescence staining with anti-α-tubulin outlined the flagellum from the base of basal body to the axonemal tip, while anti-GFP staining of eGFP-tagged IFT88 was imaged using z-stacks on a spinning-disk confocal microscope, yielding ~46 nm resolution after expansion based on the resolution calculation algorithm (Wassie et al., 2019). At the basal pool, eGFP::IFT88 consistently formed incomplete ring-like distributions around the MTDs in both non-dividing wild-type cells and zoids (Fig. 3 and Movies S2 and S3). We categorized these patterns into three categories: category 1, two distinct clusters on opposite sides of the base; category 2, one major cluster with punctate signals on the opposite side; and category 3, punctate signals encircling the entire base (Fig. 3A). To validate the results of Magnify ExM since it is not an isotropic expansion, ultrastructure expansion microscopy (U-ExM) was performed on eGFP::IFT81 cells at a lower expansion factor of 4.6x in non-dividing wild-type cells and 4.3x in zoids (Fig. S6D). U-ExM revealed more dispersed signal distributions in eGFP::IFT81 cells compared to Magnify ExM eGFP::IFT88 cells, likely reflecting protocol differences (Fig. S6A and B). Nevertheless, incomplete rings were observed consistently in 19 of 22 wild-type cells and all 15 zoids examined (Fig. S6C). This observation is different from the high-resolution imaging of IFT172 at the basal pool of flagella using 3D STORM microscopy due to the limited resolution in the z dimension to distinguish spaced trains (Araujo Alves et al., 2025). Together, these findings indicate that in *T. brucei*, IFT proteins concentrate on restricted MTDs at the flagellar base, different with *C. reinhardtii*, where IFT trains dock on all nine doublets (van den Hoek et al., 2022, Stepanek and Pigino, 2016).

**Figure 3:**
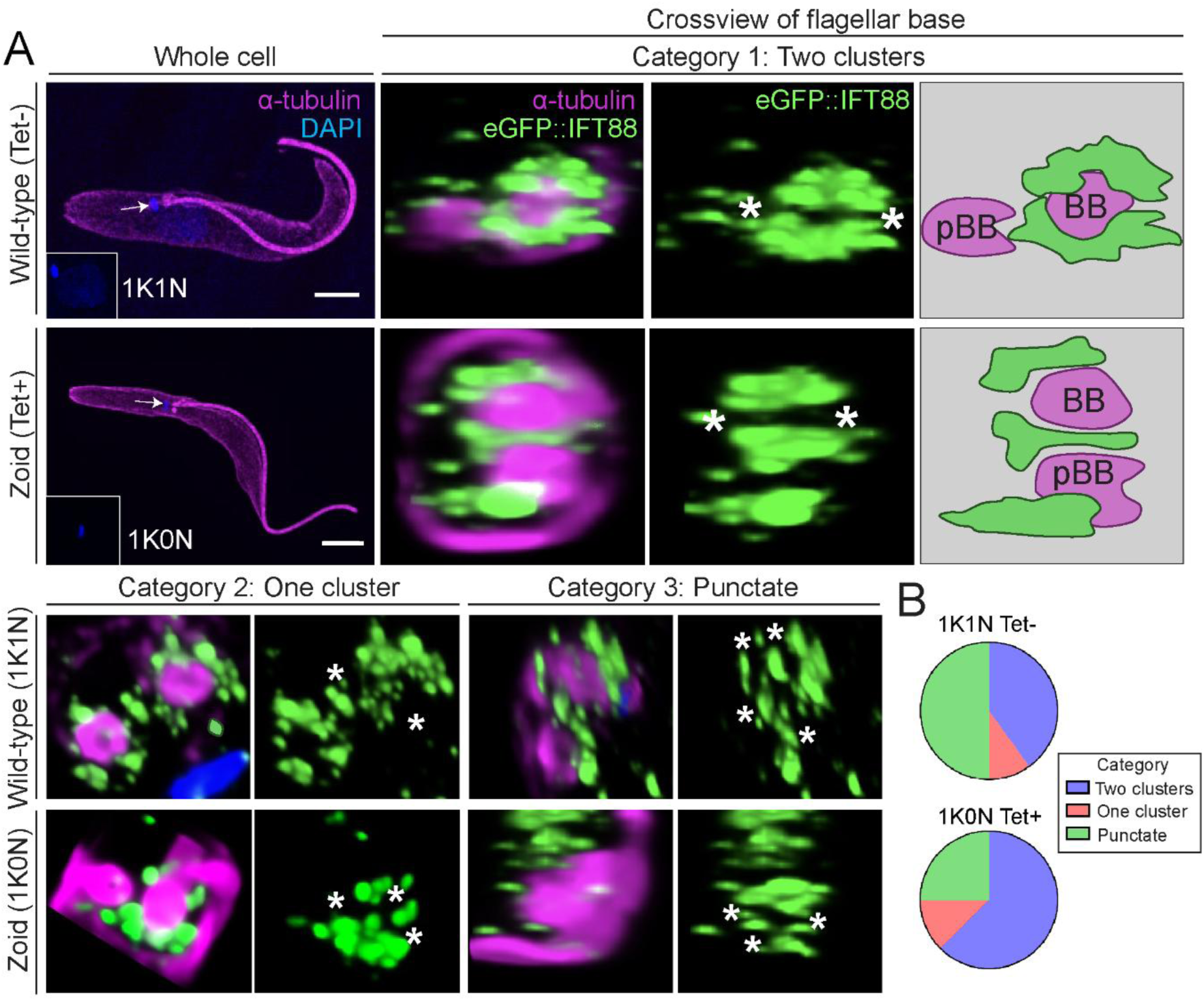
3D distribution of IFT proteins around the flagellum base compared between wild-type cells and zoids. **(A)** Magnify ExM images of eGFP::IFT88 in a non-dividing wild-type (1K1N) cell and zoid (1K0N). Three spatial categories were defined. Category 1, two separate IFT clusters on opposite sides of the base (Two clusters). Category 2, one dominant cluster with punctate IFT on the opposite side (One cluster). Category 3, punctate IFT intensities around the base (Punctate). Cartoon (right) depiction illustrating IFT distribution surrounding the basal body (BB) and the probasal body (pBB) when present in category 1. White asterisks indicate gaps between IFT clusters. **(B)** Quantification of the proportion of wild-type cells and zoids in each category for control and zoids.

### Spatial organization of IFT initial-landing trains in zoids

To further characterize the assembly and spatial organization of IFT incomplete ring formation at the flagellar base, we performed cryo-ET on *T. brucei* zoids. These cryo-tomograms revealed IFT train-like densities associated with the transition zone, similar to those IFT trains in cryo-FIB-milled wild-type cells (Fig. 4A and B and Movie S4). Measurement showed that the average train length in zoids was 268 ± 123 nm (n = 67), consistent with 265 ± 118 nm measured in in FIB-milled wild-type cells (n = 15) (Fig. S7A). To characterize the identity of these structures, small IFT density segments of these trains were computationally extracted and reconstructed by subtomogram averaging, generating an in situ cryo-EM density map at ~54.4 Å resolution (Fig. 4D and Movie S5). This map was aligned and fitted to the previously published *Chlamydomonas* IFT train structure (van den Hoek et al., 2022). While resolution was insufficient to resolve the characteristic repeating units of anterograde trains, the two major complexes, IFT-B and IFT-A, were distinguishable and arranged in a scaffold-like architecture consistent with anterograde IFT trains in *Chlamydomonas*. This structural analysis verifies that the densities observed in *T. brucei* zoids represent anterograde IFT trains.

**Figure 4:**
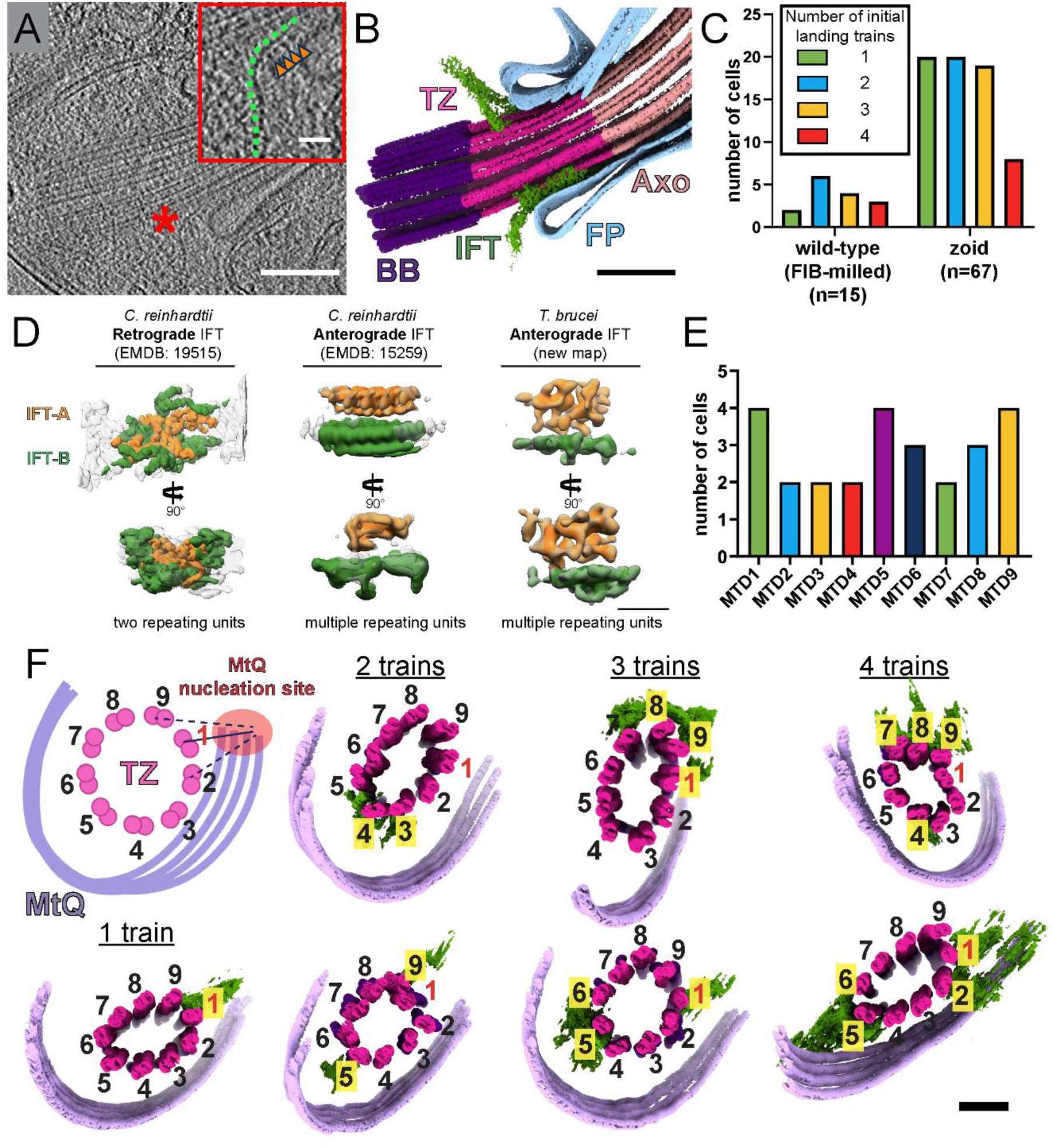
A limited number of IFT trains simultaneously land at the transition zone. **(A)** Cryo-ET tomogram slice of a zoid focused at the flagellum base, showing IFT train-like densities near the transition zone. Scale bar 200 nm. Inset, zoomed-in view of an initial landing IFT train (red asterisk), revealing distinguishable IFT-B (green dotted line) and IFT-A (orange arrows) densities. Scale bar 50 nm. **(B)** 3D annotation of the tomogram in (A), showing cytoplasmic IFT trains (green) approaching the transition zone (magenta), located distal to the basal body (BB, purple), proximal to the axoneme (Axo, pink), and within the flagellar pocket (FP, light blue). Scale bar 200 nm. **(C)** Quantification of total IFT trains at the initial landing stage in cryo-FIB-milled wild-type (n = 15) and zoid (n = 67) tomograms, showing a maximum of four trains per cell. (**D)** Averaged maps of IFT from *C. reinhardtii* show distinct configurations of the IFT-A and IFT-B complexes between retrograde (EMD-19515) and anterograde (EMD-15292). Averaging of IFT trains near the transition zone from *T. brucei* zoids at 54.4 Å showing similar architecture to anterograde trains of *Chlamydomonas*. Scale bar 20 nm. **(E)** Occupancy of initial-landing IFT trains on the nine MTDs of the transition zone, numbered relative to the MtQ nucleation point (n = 11). **(F)** Cross-sectional 3D views of transition zones (magenta) in zoids showing 1 to 4 initial-landing IFT trains (green). MTDs are numbered using the MtQ-based scheme (diagram, top left). In all cases, no more than three neighboring MTDs carried IFT trains, with additional trains positioned on the opposite side of the transition zone. Highlighted numbers in yellow indicate occupied MTDs. For the 2 or 3 train cases, examples with adjacent and bilateral train distributions are shown. For the 4 train cases, examples with 3:1 train or 2:2 distributions are shown. Scale bar 100 nm.

To further investigate whether anterograde IFT trains are restricted in the number of MTDs they access at the transition zone, we quantified the total number of IFT trains at the initial landing stage in zoids (Fig. 4C). In FIB-milled wild-type cells, we also observed between 1 and 4 trains at initial landing stage per transition zone within a cell. Though FIB-milling frequently resulted in the partial removal of the transition zone MTDs in the lateral direction, which possibly removed initial landing IFT trains, we generated three tomograms at the flagellum base in which the milling angle preserved the nine MTDs of the transition zone for the initial IFT landing (Fig. S2). We still observed less than nine trains docking, with three initial landing IFT trains in one of the cell tomograms and only one initial landing IFT train in the other two tomograms, suggesting the possibility that FIB-milling is not the cause of the 1-4 total train limit at the initial landing stage of wild-type cells. To further conclude if there are limited number of trains accessing the transition zone for landing, we examined zoids, which retain the circular arrangement of nine MTDs at the transition zone. We observed between 1 and 4 initial landing trains per zoid, suggesting that anterograde IFT trains are limited to dock to a subset of the nine MTDs at any given time. This finding is different with previous observations in *Chlamydomonas*, where anterograde IFT trains were simultaneously associated with all nine MTDs at the transition zone (van den Hoek et al., 2022, Stepanek and Pigino, 2016).

IFT train landing follows a defined spatial pattern across the MTDs of the transition zone. Consistent with *C. reinhardtii*, each train in *T. brucei* was observed to land on a single MTD at a time. However, the patterning of train distribution across MTDs in *T. brucei* varied depending on the total number of trains per cell. The axonemal MTD numbering is typically defined by the conserved positioning of the PFR between MTDs 4-7 (Lacomble et al., 2009). Because our zoid tomograms rarely captured the PFR due to the narrow imaging window required for high resolution visualization of IFT trains at the base, we established an alternative numbering system using the microtubule quartet (MtQ) as a reference point (Dong et al., 2020). The MtQ is an array of four microtubules nucleated near the basal bodies that wrap around the FP and extend along the flagellar attachment zone (Lacomble et al., 2009). The MTD closest to the MtQ nucleation site was defined as number 1, with numbering proceeding clockwise when viewed from the distal end (Fig. 4F). Based on this MtQ-based scheme, we found that all nine MTDs within the transition zone can accommodate trains at the initial landing stage (n = 11), suggesting that IFT docking is not restricted to specific doublets (Fig. 4E and F). Despite this flexibility, the trains exhibited a consistent spatial organization. No more than 3 adjacent MTDs carried trains simultaneously, and remaining trains localized to opposite sides of the transition zone. Across cells with one to four total trains, we identified seven distinct spatial configurations, including 1:0, 1:1, 2:0, 2:1, 3:0, 3:1, and 2:2, indicating distinct arrangements of trains on opposing sides. These patterns suggests that IFT organization during initial landing is not governed by specific MTD identity, but rather by spatial coordination limiting adjacent trains occupancy.

### IFT-46 depletion disrupts IFT train integrity and landing organization

To determine the role of the IFT-B complex in the formation of IFT trains at the initial landing stage, we knockdown IFT46 in zoids by dual RNAi, an IFT-B protein which was previously shown as crucial to IFT-B complex stability in *C. reinhardtii* (Hou et al., 2007). RT-PCR of Centrin4-IFT46-RNAi confirmed a decrease in expression of Centrin4 and IFT46 over a 96-hour tetracycline induction (Fig. S7C). Then, the RNAi phenotype of IFT46 was evaluated by a decrease in flagellum length following induction. Similar to our zoid tomogram analysis, we focused on the mature flagellum in non-dividing wild-type cells, which featured an average flagellum length of 15.6 ± 3.02 µm (n = 99), while the average length in zoids was 8.27 ± 4.07 µm (n = 157) after 72 hours, and 7.84 ± 4.58 µm (n = 189) after 96 hours induction (Fig. S7B).

Cryo-tomograms targeted to the mature flagellum base in IFT46 knockdown zoids enabled the visualization of IFT train-like densities (Fig. 5A and B and Movie S6). The average length of these initial landing trains in IFT46-depleted zoids was 261 ± 108 nm (n = 62), similar to that observed in FIB-milled wild-type cells with 265 ± 118 nm (n = 15) and zoids with 268 ± 123 nm in length (n = 67) (Fig. S7A). However, their widths were significantly increased, averaging 34.8 ± 8.33 nm in IFT46-depleted zoids compared to 25.3 ± 5.83 nm in wild-type and 25.6 ± 5.32 nm in zoids (Fig. 5C). These findings suggest that loss of IFT46 compromises the structural integrity of IFT trains, making them wider without affecting length. To assess whether IFT46 depletion alters the occupancy of initial landing trains at the transition zone, we first quantified the total number of trains. Compared with wild-type cells and zoids, IFT46 knockdown reduced the total number of trains, with 33 of 62 zoids tomograms containing one train, 16 tomograms containing two, and 13 tomograms containing three; none exhibited four trains, which exists in wild-type and zoids (Fig. 5E). We then evaluated occupancy across individual MTDs using the MtQ-based numbering scheme and found that, despite reduced numbers, initial landing trains still docked to all nine MTDs of the transition zone as in zoids (Fig. 5D and F). These findings indicate that IFT46 depletion disrupts overall train assembly without abolishing the intrinsic ability of IFT trains to associate to each MTD, implicating a transition zone-based regulatory mechanism in train assembly initiation rather than train docking.

**Figure 5:**
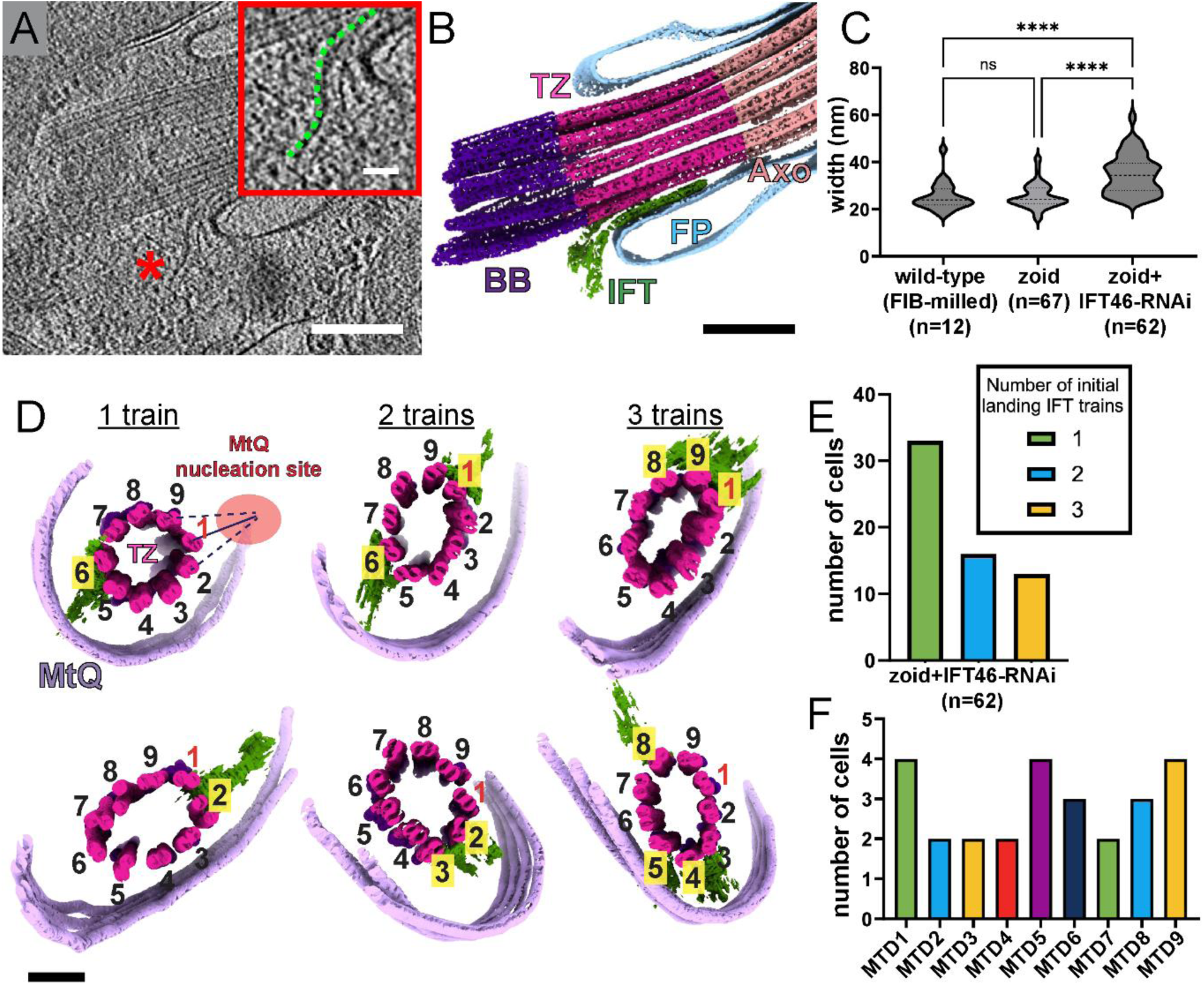
IFT46-RNAi knockdown causes reduced total number of initial landing IFT trains in zoids. **(A)** 2D tomogram slice of a IFT46-depleted zoid focused at the flagellum base. Scale bar 200 nm. Inset, zoomed-in view of an initial landing IFT train (red asterisk), revealing distinguishable IFT densities. Scale bar 50 nm. **(B)** 3D annotation of tomogram from (A) featuring trains (green) approaching the transition zone (TZ, magenta). Scale bar 200 nm. **(C)** Quantification of trains width in FIB-milled wild-type (25.3 ± 5.83 nm), zoid (25.6 ± 5.32 nm), and IFT46-depleted zoid (34.8 ± 8.33 nm) cells measured from 2D slice views. **(D)** Cross-view of 3D-annotated transition zones (magenta) with one to four total initial-landing IFT trains (green), with MTD numbering using the microtubule quartet (MtQ, lavender)-based scheme. Scale bar 100 nm. **(E)** Distribution of total initial-landing trains per cell when IFT46 is inhibited in zoids. **(F)** Distribution of initial landing IFT train occupancy at each of the nine MTDs of the transition zone numbered based on the MtQ nucleation point (n = 9).

In addition to the train assembly, we also compare the landing position of IFT trains along the transition zone in wild type cells and zoids. We quantified the relative landing position over the full length of the transition zone (D_IFT_/ D_TZ_) in FIB-milled wild-type, zoid, and IFT46-depleted zoids (Fig. S8A and B). The landing position of initial-landing trains was significantly lower in zoids and IFT46-depleted zoids compared to wild-type cells with D_IFT_/D_TZ_ ratio of 0.34 ± 0.12 (n = 67) and 0.29 ± 0.12 (n = 33), respectively, versus 0.45 ± 0.16 in wild-type cells (n = 10) (Fig. S8B). However, the average transition zone length (D_TZ_), measured from the distal end of the basal body to the basal plate, was consistent across all conditions, with 345 ± 46.4 nm in wild-type (n *=* 16), 347 ± 34.2 nm in zoids (n = 98), and 351 ± 31.3 nm in IFT46-depleted zoids (n = 93) (Fig. S8A), consistent with previously reported lengths from 200 to 400 nm (Trépout et al., 2018). Correspondingly, the placement of the FP membrane along the transition zone was also shifted downward in both zoid conditions. The distance of the FP base to the distal end of the basal body (D_FP_) over the full length of transition zone, D_FP_/D_TZ_, was 0.31 ± 0.11 in zoid (n = 98), 0.29 ± 0.15 in IFT46-depleted zoids (n = 53), significantly lower than 0.71 ± 0.19 in wild-type cells (n = 16) (Fig. S8C). This correlation between the FP membrane position and IFT landing suggests that structural elements associated with the FP membrane could play a role in IFT engagement at the transition zone.

## Discussion

In this study, we used cryo-FIB milling and cryo-ET to define how IFT proteins initiate entry into the flagellum of *T. brucei* at nanometer resolution, revealing previously unrecognized features of the IFT train landing process. By establishing enucleated zoids as a proxy for wild-type cells, we were able to reproducibly visualize the entire transition zone and map the spatial organization of individual IFT trains in situ. Complementary Magnify ExM provided crucial cellular context by revealing discrete gaps within the basal IFT pool, enabling direct comparison between wild-type and zoid architectures. Together, these approaches demonstrated that *T. brucei* does not assemble IFT trains simultaneously on all nine MTDs at the transition zone, in contrast to the full MTD occupancy reported in *Chlamydomonas* and to ring-like distributions inferred from super-resolution fluorescence imaging in *T. brucei* (Araujo Alves et al., 2025, van den Hoek et al., 2022). Instead, our cryo-ET analyses, supported by ExM characterization, revealed a spatially asymmetric and alternating pattern of IFT train landing. Integrating these high-resolution cryo-ET and expanded-cell imaging strategies allowed us to uncover how the geometry of the transition zone, the MtQ, and local membrane architecture collectively shape the landscape of IFT entry. These findings point to a previously unrecognized regulatory mechanism that spatially constrains IFT landing at the transition zone before doublet selectivity and anterograde trafficking are established within the axoneme.

To determine how this spatial restriction is established, we developed an MtQ-based numbering system (Dong et al., 2020, Gheiratmand et al., 2013). We found that all nine MTDs are competent for IFT landing, indicating that doublet selectivity arises only after complete landing. However, no more than three adjacent MTDs host trains at once, with additional trains positioned on the opposite side of the transition zone, revealing a previously unrecognized regulatory mechanism that limits simultaneous occupancy across the entire circumference. We next used this strategy to examine the role of the IFT-B complex by depletion of IFT46. Although train formation and initial landing still occurred, fewer were assembled overall, and those that did were significantly wider, revealing compromised structural integrity. These findings suggest that initial landing is not strictly dependent on intact IFT-B scaffolding, but that IFT-B integrity is required to preserve train architecture. Together, these results establish that our new strategy uncovered an early-stage gating mechanism at the flagellar base that restricts how many IFT trains may enter the transition zone at once. Importantly, this regulatory step appears to operate independently of the IFT-B complex integrity, pointing to additional factors that govern IFT entry and spatial organization before doublet-specific trafficking is imposed in the axoneme.

The use of enucleated zoids was critical to uncovering this regulatory landscape. Zoids maintain motility but have reduced cytoplasmic volume due to lack of nucleus. This simplification makes them exceptionally well suited for whole-cell cryo-ET without the need for cryo-FIB milling (Sun et al., 2018, Bregy et al., 2023). Because zoids are non-dividing *T. brucei*, they provide a synchronized population with fully mature flagella, eliminating variability from cell-cycle–dependent flagellar growth. Importantly, IFT train size and morphology in zoids closely matched those observed in cryo-FIB–milled wild-type cells, demonstrating that the absence of the nucleus does not disrupt IFT train architecture. These cryo-ET observations were further supported by two ExM approaches, including Magnify ExM, which establishes a platform for future visualization of multiple IFT-A and IFT-B subunits, motors, and transition zone components within the same expanded flagellar base (Klimas et al., 2023, Cheng et al., 2023). Using this integrated strategy, we identified a maximum of four simultaneously docking trains per cell (Fig. 6), rather than occupancy of all nine MTDs as seen in *Chlamydomonas*, bridging the knowledge gap in our understanding of the IFT initiation process in *T. brucei* (van den Hoek et al., 2022, Jordan and Pigino, 2021). This difference likely reflects species-specific dynamics of flagellar assembly. *Chlamydomonas* flagella undergo continuous assembly and disassembly, requiring constant delivery or dispatching of precursors, whereas the mature *T. brucei* flagella, like photoreceptors or spermatozoa, are structurally stable and no longer elongate once completed (Marshall and Rosenbaum, 2001, Fort et al., 2016, Bertiaux et al., 2018b). Consistent with this, prior studies show that IFT levels are elevated during elongation compared to in mature *T. brucei* flagella (Bertiaux et al., 2018b). However, a limitation of the zoid system is that it restricts analysis to mature flagella, preventing direct assessment of whether early assembly stages exhibit different IFT landing patterns. Moreover, the limited field of view inherent to cryo-FIB milling rarely captures duplicated flagella in dividing wild-type cells, complicating stage-specific comparisons.

**Figure 6:**
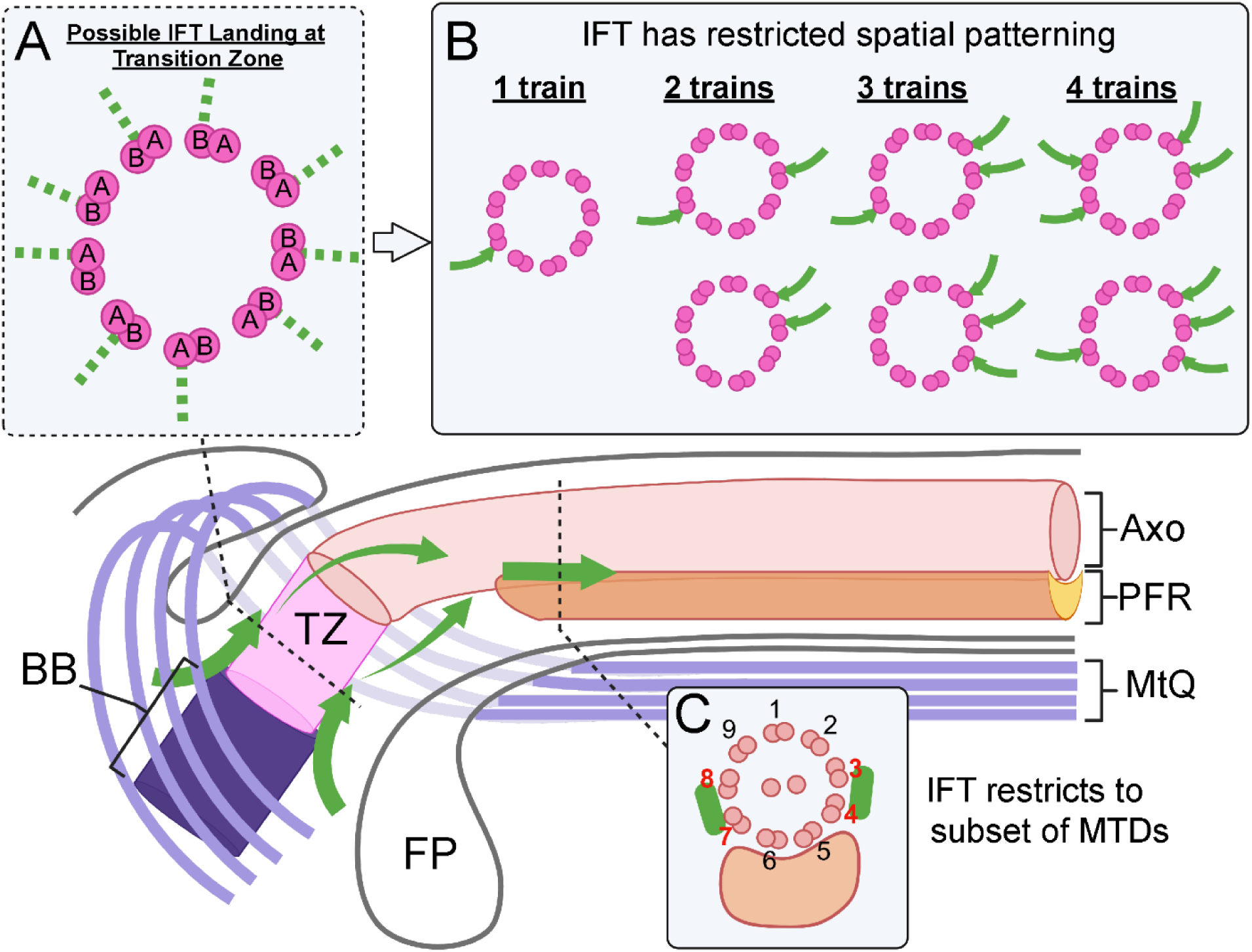
Model of spatial organization of anterograde IFT initial-landing at transition zone in *T. brucei*. **(A)** Schematic illustration showing that all nine MTDs (magenta) within the transition zone (TZ) serve as potential landing sites for anterograde IFT trains at the initial landing stage (green dashed lines). **(B)** However, a maximum number of four trains are observed simultaneously at the transition zone. No more than three adjacent MTDs host trains at once, with the remaining trains positioned on opposite sides forming two spatially separated groups. **(C)** After complete docking, trains progress from the transition zone into the proximal axoneme (Axo), and then enter the region where the axoneme is associated with the PFR region, at which point they become restricted to MTDs 3-4 and 7-8, consistent with previously described doublet-selective IFT trafficking in *T. brucei* (Bertiaux et al., 2018a, Araujo Alves et al., 2025).

Our cryo-ET analyses of zoids also revealed a previously unrecognized positional shift in IFT docking along the transition zone. Although train size and morphology were comparable between wild-type cells and zoids, initial IFT landing occurred consistently lower along the transition zone in zoids, an effect that persisted upon IFT46 depletion. This observation suggests that docking position is regulated by cellular architecture rather than by IFT complex composition alone. A likely contributor is Centrin4, which localizes to the bi-lobe encircling the neck of the FP and is tightly connected to the FP collar (Shi et al., 2008). Recent work has shown that this structure is essential for the de novo biogenesis of the FP and its collar (Zelená et al., 2025, Morriswood et al., 2013). Consistent with this role, both zoids and IFT46-depleted zoids exhibited a lower placement of the FP membrane along the transition zone relative to wild-type cells. The correlation between FP membrane positioning and anterograde IFT docking height suggests that membrane geometry or FP-associated proteins may play an active role in IFT entry and spatial organization at the flagellar base, revealing a new control distinct from axonemal doublet selection. Future studies combining molecular perturbation of FP components with in situ structural analyses will be essential to dissect how FP architecture and membrane-associated factors and collar architecture contribute to IFT regulation at the basal pool in *T. brucei*.

Although mature *T. brucei* flagella no longer elongate, they continue to support IFT entry. Recent work has shown that distal remodeling events such as radial spoke turnover at the flagellar tip (Abbühl et al., 2025), remain active in fully constructed flagella, providing a biological rationale for this continued IFT activity. At the same time, these observations highlight the need for mechanisms that tightly regulate where and when trains enter the flagellum. Consistent with this idea, increasing IFT rate in a “locked” mature flagellum does not trigger additional elongation (Bertiaux et al., 2018b), suggesting that IFT entry is spatially gated for growth control. In addition, the existence of a basal pool of IFT proteins has been well recognized in our studies, but how components are delivered to the basal pool prior to train assembly remains unclear. One possibility is active transport along cytoskeletal microtubule tracks, as shown for the ciliary protein CCDC66 trafficking to the basal bodies in mammalian cells (Conkar et al., 2019). In *T. brucei*, the kinesin motor Kin2a was previously identified as crucial for flagellar biogenesis and was also localized along the FAZ, which includes the MtQ, raising the potential role of the MtQ as a delivery route to the basal pool (Douglas et al., 2020). Determining whether Kin2a trafficking within the FAZ-MtQ contributes to IFT pool assembly will require future investigation.

We identified the cryo-ET densities as anterograde IFT based on periodicity measurements of 2D images and averaged scaffold structures resembling IFT-A and IFT-B complexes from *Chlamydomonas* anterograde trains (Lacey et al., 2023). However, we did not detect densities characteristic of retrograde IFT, which display a ~45 nm periodicity following structural rearrangement at the flagellar tip in *C. reinhardtii* (Lacey et al., 2024). This absence is consistent with the lack of retrograde trains at the ciliary base in cryo-FIB-milled *Chlamydomonas* (van den Hoek et al., 2022). One hypothesis is that retrograde trains disassemble before reaching the proximal flagellum, with their components diffusing back into the basal pool rather than remaining as intact assemblies. Notably, we find that anterograde IFT trains in *T. brucei* can associate to the A-tubule, B-tubule, or the inter-tubule interface at the flagellar base. This contrasts with *C. reinhardtii*, where anterograde trains are confined to B-tubules, and retrograde trains to A-tubules (Stepanek and Pigino, 2016), a spatial segregation thought to prevent collisions and enable simultaneous bidirectional transport along a single doublet. The absence of such tubule-specific segregation in *T. brucei* raises the question of how bidirectional IFT is coordinated. One assumption is that retrograde transport preferentially occurs on MTDs that are transiently unoccupied by anterograde trains when reaching to transition zone. Such a mechanism could account for both the limited number of anterograde trains at initial landing and the absence of retrograde assemblies at the flagellar base. Resolving this coordination will require studies of the axoneme region between the transition zone and the onset of PFR initiation region. Future application of correlative light and electron microscopy as well as cryo-ET with molecular feature markers will be critical to localize retrograde complexes in situ and to define the structural transitions that accompany their return. Together, these approaches may uncover new principles governing bidirectional IFT organization and regulation in cilia or flagella.

## Methods

### Cell Culture and Transfection

Procyclic form *T. brucei* from the double-inducible YTaT1.1 (DIY) or 29.13 cell lines engineered for tetracycline-inducible RNAi were cultured in Beck’s medium (Cytiva) supplemented with 10% heat-inactivated Tet-free fetal bovine serum (BioTechne) and 10 µg/ml Gentamicin (Gibco, Cat# 15-710-064) at 28°C and 5% CO_2_. 29.13 cell growth medium included 10 µg/ml Geneticin (Gibco, Cat# 10-131-035) and 10 µg/ml Hygromicin B (ThermoFisher Scientific) (Ruben et al., 1983, Li et al., 2017, Wirtz et al., 1999). YTaT1.1 cell growth medium included 5 µg/ml puromycin. RNAi knockdown of Centrin4 was performed in 29.13 cells as described previously (Shi et al., 2008, Sun et al., 2018). For single or combined RNAi knockdown of Centrin4 and IFT46, a 495-bp fragment (nucleotides 311-806) of the IFT46 gene was cloned into the p2T7-Centrin4 plasmid via XbaI digestion, yielding the p2T7-Centrin4 and p2T7-Centrin4-IFT46 plasmid. For transfection, 15 µg of the p2T7-Centrin4-IFT46 plasmid was linearized by NotI (NEB) digestion, followed by electroporation into ~5 x 10^7^ DIY cells, and serial dilution with 10 µg/ml phleomycin (Millipore Sigma) to generate a stable cell line.

For endogenous N-terminal tagging, the pPOTv7 tagging system was used (Dean et al., 2015, Paterou et al., 2025). Each protein was tagged using a forward primer composed of the last 70 base pairs of the 5’ untranslated region and 20 base pairs annealing to the pPOTv7-eGFP-blast plasmid, and a reverse primer composed of 18 base pairs annealing to the GS-linker of the plasmid and the first 72 base pairs of the open reading frame of the gene.

### Isolation of zoids

Centrin4-RNAi or dual Centrin4-IFT46-RNAi cells were cultured with 10 µg/ml tetracycline for 72 hours to generate zoids or IFT46-depleted zoids, respectively. Zoid isolation was performed as described previously (Sun et al., 2018). 3-5 x 10^7^ cells were harvested and pelleted by centrifugation at 3000 rpm for 7 min and resuspended in 1 ml medium. Enucleate zoids were enriched over three rounds of low-speed centrifugation using a swing-bucket rotor centrifuge (Eppendorf 5427R, Rotor S-24-11-AT), retaining cell culture from the top portion of the supernatant containing zoids after each round. The centrifugation rounds were performed as follows: (i) 500 rpm for 2 min, 800 rpm for 2 min, and 1400 rpm for 5 min, retaining 70% of the supernatant volume, (ii) 800 rpm for 3 min and 1500 rpm for 5 min, retaining 70% of the supernatant volume, and (iii) 1800 rpm for 3 min, retaining 60% of the supernatant volume, which was the most enriched fraction containing zoid or IFT46-depleted zoid cells.

### Immunofluorescence staining and light microscopy

Cells were pelleted by centrifugation at 4000 rpm for 2 min, washed and resuspended in 1x PBS (pH 7.2), and settled onto coverslips for 30 min, followed by fixation in methanol at −20°C for 5 min. Following fixation, coverslips were washed with 1x PBS for 10 min and blocked with 3% BSA in 1x PBS for 1 hour at room temperature or overnight at 4°C. The anti-α-tubulin antibody was used to stain the cell body and the anti-PFR2 antibody was used to stain the flagella (see Table 1). Nuclear and kinetoplast DNA were stained using 5 μg/ml DAPI (D1306; Invitrogen). Z-stacks for autofluorescence intensity measurements were acquired using an A1 confocal microscope (Nikon) with a 60x (NA 1.4) oil objective. Fluorescent images were processed and analyzed using FIJI (Schindelin et al., 2012).

### Western Blot

Cells were pelleted by centrifugation at 4000 rpm for 2 min, washed once with 1x PBS (pH 7.2), resuspended in Laemmli SDS 4x sample buffer (ThermoFisher Scientific, Cat# J60015.AC), and incubated at 100°C for 5 min. Protein sample equivalent to 3×10^6^ total cells for eGFP::IFT81/88 staining and, due to the significantly higher expression of tubulin, 1.5×10^6^ total cells for α-tubulin staining were run on separate 10% SDS-PAGE gels in 1x Tris/Glycine/SDS buffer (Bio-rad, Cat# 1610772) (90-150V, 1.5 hours). Gels were transferred onto Immuno-blot® PVDF membranes (Bio-Rad, Cat# 1620177) in a wet-blotting chamber in 1x Tris/Glycine buffer (Bio-rad, Cat# 1610771) (0.4 A, 1.5 hours). Membranes were blocked in 5% dry milk dissolved in 1x Tris-buffered saline (pH 8.0) and 0.1% Tween (1x TBST) for 1 hour at room temperature, followed by overnight primary antibody staining (see Table 1). Membranes were washed with 5% milk in 1x TBST 3x, followed by secondary antibody staining (see Table 1) for 3 hours at room temperature. Western blots were imaged using a LI-COR Odyssey CLx imager.

### RT-PCR

Total RNA was isolated from 8×10^7^ cells using TRIzol® reagent (Invitrogen, USA, Cat# 15596018). Isolated RNA was treated with RNase-free DNase I (Invitrogen, USA, Cat# 18068015) and the first-strand cDNA was synthesized using M-MLV reverse transcriptase (Invitrogen, USA) with oligo(dT)12–18 primer (Fermentas, USA). The RNA yield from samples of various induction times were measured by Nanodrop and normalized after DNase I treatment, before first-strand synthesis. The cDNA level of Centrin4 and IFT46 were then monitored by PCR amplification of a 420 and 496 base-pair fragment, respectively, using primers 5’-GAACAGATCCGTGAAGCG-3’, 5’-CATCTGCATCATGACGCTC-3’, 5’-TGATGAAGTGGTTCCGGTGG-3’ and 5’-GAGGTCGCCAATGGAAGGAAC-3’. As a loading control, the cDNA level of α-tubulin was monitored by PCR amplification of a 164 base pair fragment with primers 5’-AAGCGCGCCTTCGTGCACTG-3’ and 5’-CGGGATCCCTAGTACTCCTCCACATCCTCC-3’. PCR products were imaged with a Fotodyne 60-7020 Electronic Gel Imager (Fotodyne, USA).

### Expansion Microscopy

For Magnify ExM, 1 ml of uninduced cells at 2-5 x 10^6^ cells/ml or zoids at ~1 x 10^6^ cells/ml isolated after 72-hour tetracycline induction were pelleted by centrifugation at 4000 rpm for 2 min, washed, and resuspended in 1x PBS, and settled onto coverslips for 30 min, followed by fixation with 4% PFA in 1x PBS for 20 min and washing in 1x PBS for 20-30 min. Gelation and expansion followed the Magnify protocol, which employs methacrolein-based in situ anchoring and a reinforced hydrogel. Homogenization used a denaturing buffer per protocol, followed by complete SDS removal before staining to ensure antibody accessibility (Klimas et al., 2023). For U-ExM, cells were directly fixed in growth medium with 4% PFA for 20 min at 28°C, followed by washing with 1x PBS. U-ExM gelation and expansion were then performed as described previously (Liffner et al., 2023, Gambarotto et al., 2019). For both Magnify and U-ExM, gels were stained with anti-GFP and anti-α-tubulin primary antibodies (see Table 1).

Imaging used a Nikon Eclipse Ti2 with a CSU-W1 spinning disk module and Andor Zyla 4.2 sCMOS, 60× water-immersion objective; effective lateral resolution after expansion was ~46 nm (per resolution-after-expansion calculation). Magnify expansion gel imaging was performed using an Eclipse Ti2 epifluorescence microscope (Nikon) equipped with a CSU-W1 spinning disk confocal module and an Andor v.4.2 Zyla sCMOS camera using a 60x water objective. U-ExM gel imaging was performed using an Airyscan 2 detection LSM900 microscope with 63X oil objective. Linear expansion was measured by axoneme diameters.

### Cell vitrification

Quantifoil Cu 200 mesh grids with a hole size of R2/1 for wild-type cells or R3.5/1 for zoid cells were glow discharged at 25 mA for 1 min using an EmiTech K100X glow discharger (EMS). Grids were then mounted on a Vitrobot Mk IV plunge freezer (ThermoFisher Scientific - TFS, Waltham, MA, USA), blotted from the back side using Whatman 541 filter paper (Cytiva, Cat# 1541-047), and plunged into a 1:1 mixture of liquid ethane and propane at near–liquid nitrogen temperature. ~1×10^7^ cells/ml wild-type cells and 3-5×10^6^ cells/ml isolated zoids mixed with concentrated 15 nm BSA Gold Tracer (EMS) at 1:1 (v/v) ratio were applied to grids in 4 µl volumes for plunge-freezing and manually blotted for 3-4 seconds. Plunge-frozen grids were clipped using EM auto-rings (EMS) and stored in liquid nitrogen until cryo-FIB-milling or cryo-ET acquisition.

### Cryo-FIB-milling

Cryo-FIB-milling was performed using an Aquilos 2 (TFS), a cryo-dedicated dual-beam microscope. Cryo-fixed wild-type cells on grids were sputter coated with a layer of organic platinum for 10 seconds, followed by a ~500 nm thick deposition of organic platinum applied by the gas injection system, and a final sputter coating for 15 seconds. Cell clusters were milled using a gallium ion beam to generate lamella of ~120-200 nm thickness using 0.1-0.3 nA ion beam current for rough milling and 30-50 pA for final polishing.

### Cryo-electron tomography

Cryo-FIB-milled lamella of wild-type cells or zoids were imaged using a Titan Krios electron microscope (TFS) equipped with a 300 kV field emission gun, a Selectris energy filter, and Falcon 4i direct electron detector (TFS). Tilt series were collected using Tomo5 software (TFS) at 42,000x magnification corresponding to a pixel size of 2.9 Å for wild-type lamella or at 33,000x magnification corresponding to a pixel size of 3.6 Å for zoids, and the energy filter at 20.0 eV slit width. Tilt series were collected with bidirectional or dose-symmetric acquisition schemes, each from −60° to 60° with 3° increments between tilt images. The target defocus was set to −3 to −6 μm for lamella and −5 to −10 μm for zoids. Total exposure for each tilt series was limited to 100 to 120 e^-^/Å^2^. For implementation, tilt series were collected at −1 µm defocus if volta phase plate was applied.

Motion correction of tilt images was performed using MotionCor2 (Zheng et al., 2017) in the Relion 3.1 (Zivanov et al., 2020) suite using 5×5 patches. Tomograms were aligned and reconstructed using the EMAN2 automated pipeline (Chen et al., 2019). IFT train densities were segmented using convolutional neural network training in EMAN2 (Chen et al., 2017). Microtubule doublets/triplets/quartet were manually annotated using 3dmod in IMOD 4.8.50 (Mastronarde, 1997, Kremer et al., 1996). Tomogram feature visualization and analysis, including length and width measurements and counting total IFT trains, was performed in ChimeraX (Goddard et al., 2018, Pettersen et al., 2021).

Subtomogram averaging of initial-landing IFT trains, transition zone, and axoneme was performed in EMAN2. For IFT, particles were manually picked by centering boxes on repeating IFT-A units distanced ~11.5 nm apart along the tail portion of initial landing IFT trains in zoid tomograms. A total of 816 particles from 47 tomograms were extracted using a box size of 256, binned 2x, and used to generate *de novo* initial models. For the transition zone/axoneme MTDs, particles were extracted from manually traced curves along each MTD using a 240 box size with a specified overlap of 90% between particles, binned 2x, and used to generate *de novo* initial models. For the transition zone, 2155 particles from 11 tomograms were used. For the axoneme, 2431 particles from 11 tomograms were used. Particle alignment and averaging were performed in the new 3D refinement pipeline of EMAN2.

### Statistical analysis

Statistical analyses were performed in GraphPad Prism v10. T-tests were conducted for autofluorescence intensity. Anova tests were used for all other analyses. In the graphs, * indicates p < 0.05, ** indicates p < 0.01, **** indicates p < 0.0001, and not significant (ns) indicates p > 0.05.

## Data Availability

Electron tomography data will be made available on the EMDB.

## Acknowledgements

We thank James Conway and the Pittsburgh Center for CryoEM (RRID:SCR_025216) for training and support of data collection. This project was supported, in part, by the University of Pittsburgh, the School of Medicine, the Department of Structural Biology, and the NIH (grants S10-OD-019995 and S10-OD-025009). A portion of this research was supported by the NIH (grants U24 GM139168 and U24 GM139166) and performed at the Midwest Center for Cryo-Electron Tomography (MCCET) and the Cryo-EM Research Center in the Department of Biochemistry at the University of Wisconsin-Madison and the Stanford-SLAC Cryo-ET Specimen Preparation Center (SCSC) at Stanford-SLAC. A portion of this work was supported by the NIH (grant R01GM150905). The content is solely the responsibility of the authors and does not necessarily represent the official views of the National Institutes of Health. We thank Katherine Helfrich, Mike Calderon, and the University of Pittsburgh Center of Biologic Imaging (CBI) for training and equipment access. We thank Maisie Weiss for her assistance with data visualization. We thank Cynthia He (National University of Singapore) for providing the α-YFP antibody. We thank Jonathan Coleman for providing access and assistance with the LI-COR scanner.

