## Supplemental Figures for "In situ cryo-ET reveals restricted docking of intraflagellar transport at the base of the trypanosome flagellum"

| Primary antibody | Raised in | Antibody source (Cat#) | Usage | Antibody dilution |
|---|---|---|---|---|
| Anti-alpha-tubulin (Clone B-5-1-2) | Mouse | Abcam (ab11304) (IF/WB) | IF/WB | 1:2000 (IF)<br>1:5000 (WB) |
| | | ThermoFisher (32-2500) (U-ExM) | U-ExM | 1:150 |
| Anti-beta-tubulin | Mouse | Proteintech (66240-1) | Magnify ExM | 1:2000 |
| Anti-GFP | Chicken | Abcam (ab13970) | Magnify ExM | 1:1000 |
| | Rabbit | Fisher Scientific (NC9589665) | U-ExM | 1:2000 |
| Anti-PFR2 | Rabbit | Gifted by Cynthia He (NUS) | IF | 1:500 |
| Anti-YFP | Rabbit | Gifted by Cynthia He (NUS) | WB | 1:1000 |
| **Secondary antibody** | **Raised in** | **Antibody source (Cat#)** | **Usage** | **Antibody dilution** |
| Alexa Fluor® 488 anti-chicken IgY | Goat | Jackson (103-547-008) | Magnify ExM | 1:1000 |
| Alexa Fluor™ 488 anti-rabbit IgG (H+L) | Goat | Invitrogen (A11008) | IF | 1:1000 |
| | | Invitrogen (A11034) | U-ExM | 1:500 |
| Alexa Fluor™ 568 anti-mouse IgG (H+L) | Goat | Invitrogen (A11004) | IF | 1:1000 |
| Alexa Fluor™ 555 anti-mouse IgG (H+L) | Goat | Invitrogen (A21422) | U-ExM | 1:500 |
| CF® 640 anti-mouse IgG (H+L) | Donkey | Biotium (20177) | Magnify ExM | 1:1000 |
| IRDye® 800CW anti-rabbit | Goat | LI-COR (926-32211) | WB | 1:5000 |
| IRDye® 680LT anti-mouse | Goat | LI-COR (926-68020) | WB | 1:20000 |

**Table 1:** All primary and secondary antibodies applied in this study.

A

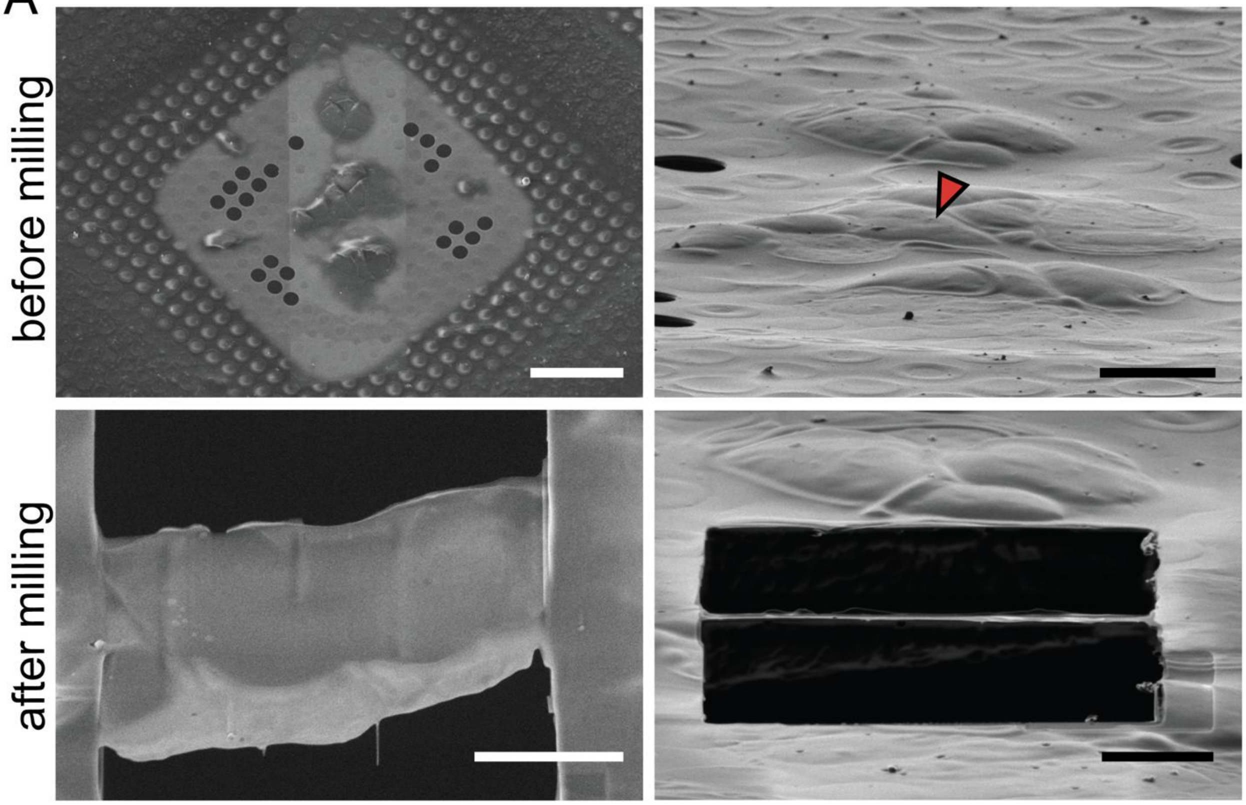


B

| Milling session time (hours) | Number of lamella generated | Number of tomograms at flagellum base | Number of tomograms with visible IFT train density | Number of tomograms with intact 9+0 transition zone |
|---|---|---|---|---|
| 9 days | 66 | 54 | 21 | 3 |
| | 7-8/per day | ~30% | ~15% | 5% |

**Supplemental Figure 1: Cryo-FIB-milling of *T. brucei* frequently removes the 9+0 transition zone cylinder. (A)** Cryo-focused ion beam (cryo-FIB) milling workflow showing targeted lamella preparation at the flagellar base (red arrow). (**B)** Quantification of all 66 lamellae generated, including 21 out of 54 tomograms containing the flagellar base with the high-quality images to visualize IFT train densities, and only 3 tomograms (5% in total number of tomograms) which preserve all nine MTDs of the transition zone.

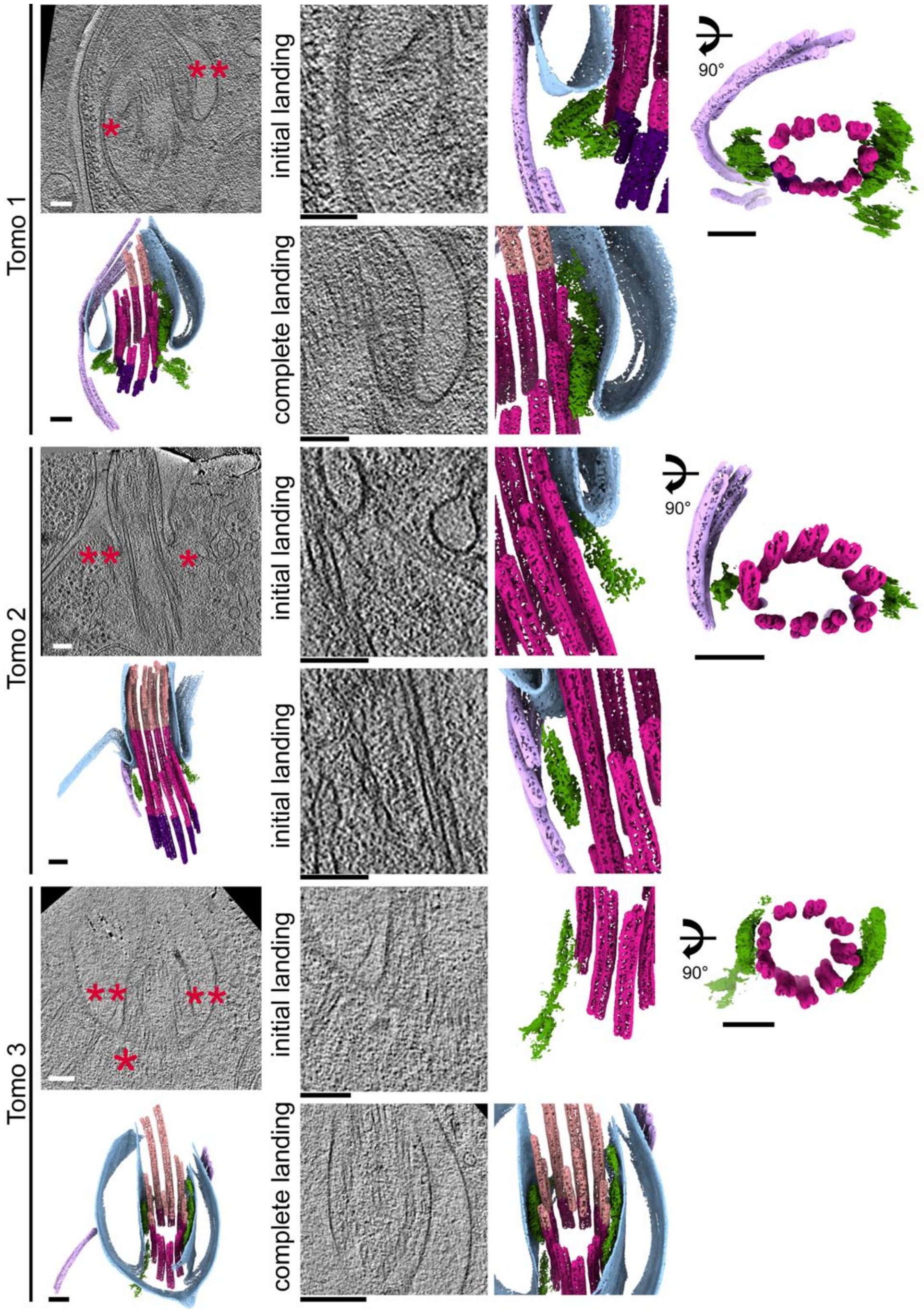


**Supplemental Figure 2: Wild-type FIB-milled cells preserve all nine MTDs of transition zone and exhibit restricted number of IFT initial landing trains at base.** Among 54 tomograms of cryo-FIB-milled cells, three retained all nine MTDs of the transition zone at the flagellar base. The two stages of landing trains were observed, including the complete landing trains (double red asterisks). Either one or two initial landing (single red asterisk) were observed, rather than nine trains surrounding all nine MTDs. These results indicate that the limited number of initial landing trains is not an artifact of FIB-milling but suggests an intrinsic restriction at base. Scale bars 100 nm.

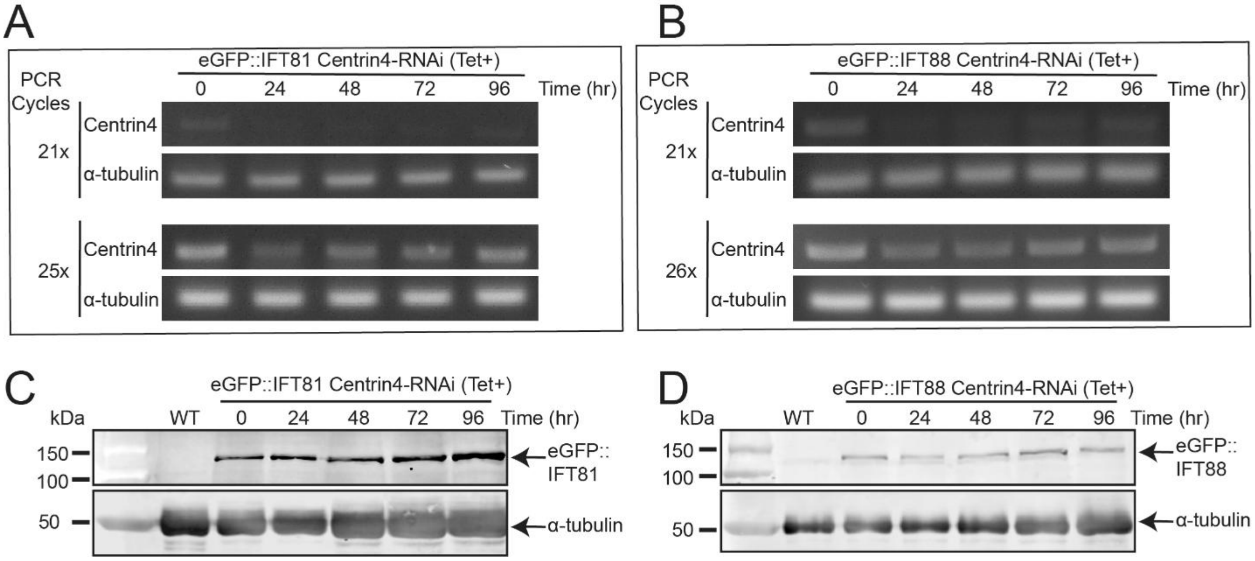


**Supplemental Figure 3. The expression of eGFP-tagged IFT81 and IFT88 in Centrin4 knockdown cells. (A-B)** RT-qPCR analysis confirming efficient Centrin4 knockdown in cells expressing eGFP::IFT81 or eGFP::IFT88 after 96 hours of tetracycline-induced RNAi. **(C-D)** Western blot analysis showing that endogenous expression of eGFP::IFT81 and eGFP::IFT88 are maintained in Centrin4 RNAi cells following 96 hours of tetracycline-induction.

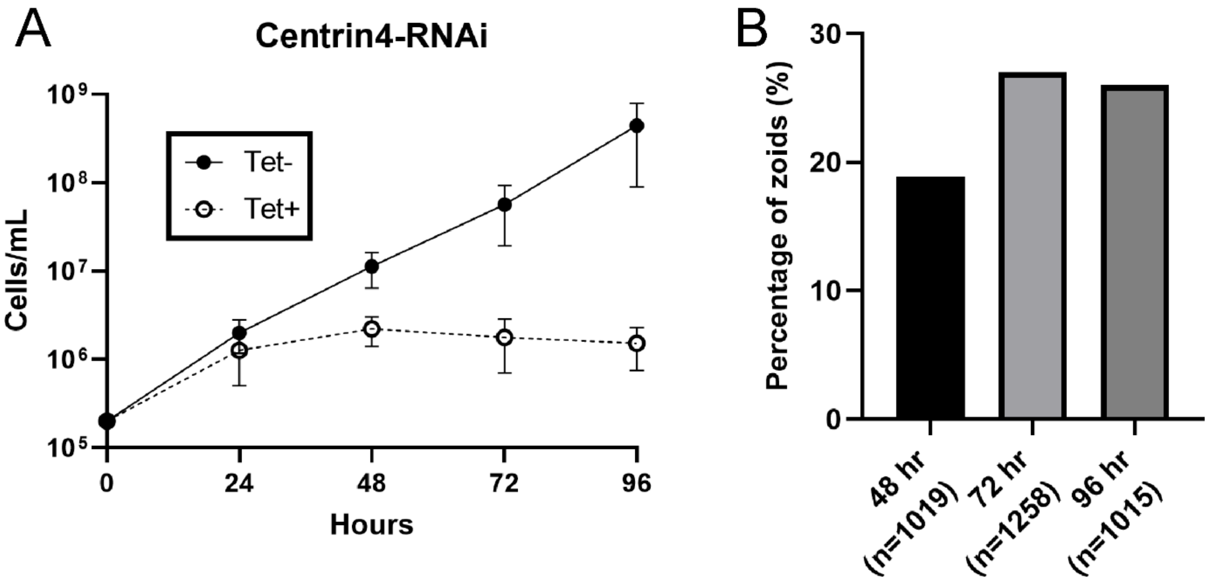


**Supplemental Figure 4: Characterization of zoids generation by Centrin4-RNAi induction. (A)** Cell growth was inhibited after 48 hours of Centrin4 knockdown by tetracycline-induced RNAi. **(B)** Quantification of zoid generation indicated the highest percentage was 27% at 72 hours post-induction, compared to 19% at 48 hours, and 26% at 96 hours.

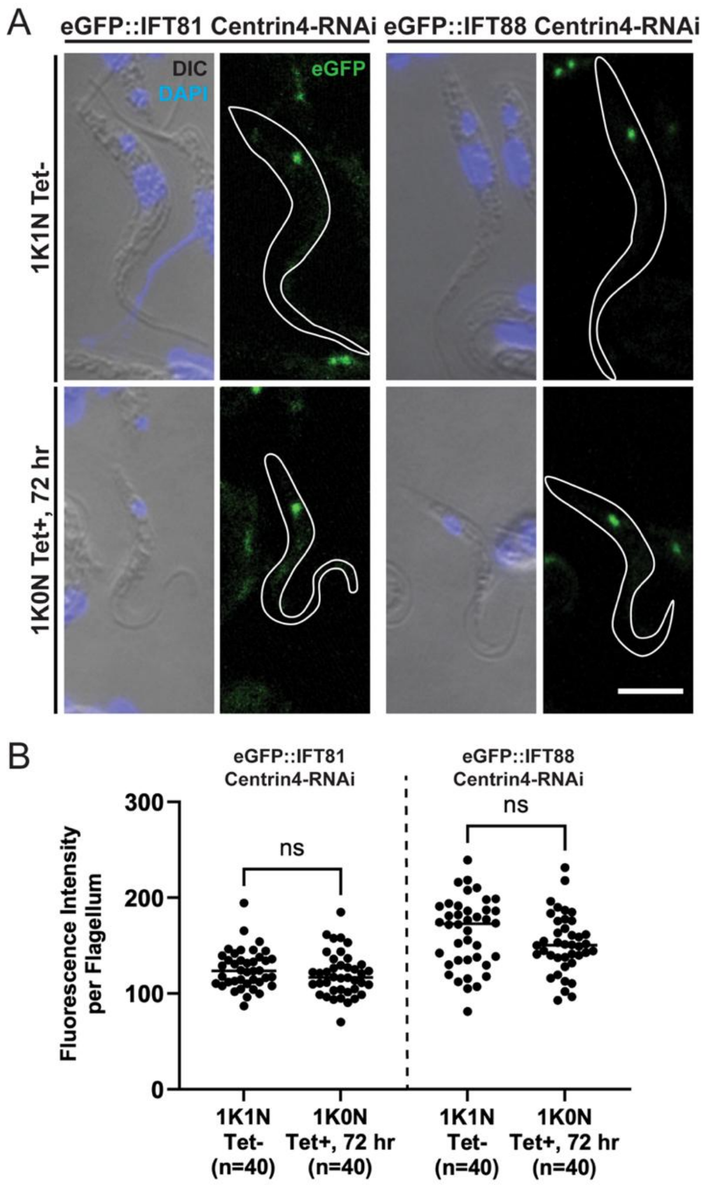


**Supplemental Figure 5: Quantification of eGFP::IFT81 and eGFP::IFT88 concentration at the flagellum base shows no difference between wild-type cells and zoids. (A)** Centrin4-RNAi cells endogenously expressing eGFP::IFT81 or eGFP::IFT88 were imaged either in non-dividing wild-type cells (1K1N shown by DAPI) or in zoids (1K0N), after 72 hours of tetracycline induction. Scale bar 5 μm. **(B)** Quantification of autofluorescence intensity at the flagellar base (arbitrary fluorescence units, A.F.U.) showed comparable levels of eGFP::IFT81 in wild type cells (125.4 ± 20.50 A.F.U.) and zoids (119.7 ± 22.84 A.F.U.), as well as similar eGFP::IFT88 levels between wild type (162.4 ± 36.32 A.F.U.), and zoids (150.7 ± 30.42 A.F.U.). These measurements indicate that Centrin4 depletion does not change the concentration of IFT proteins at the flagellar base.

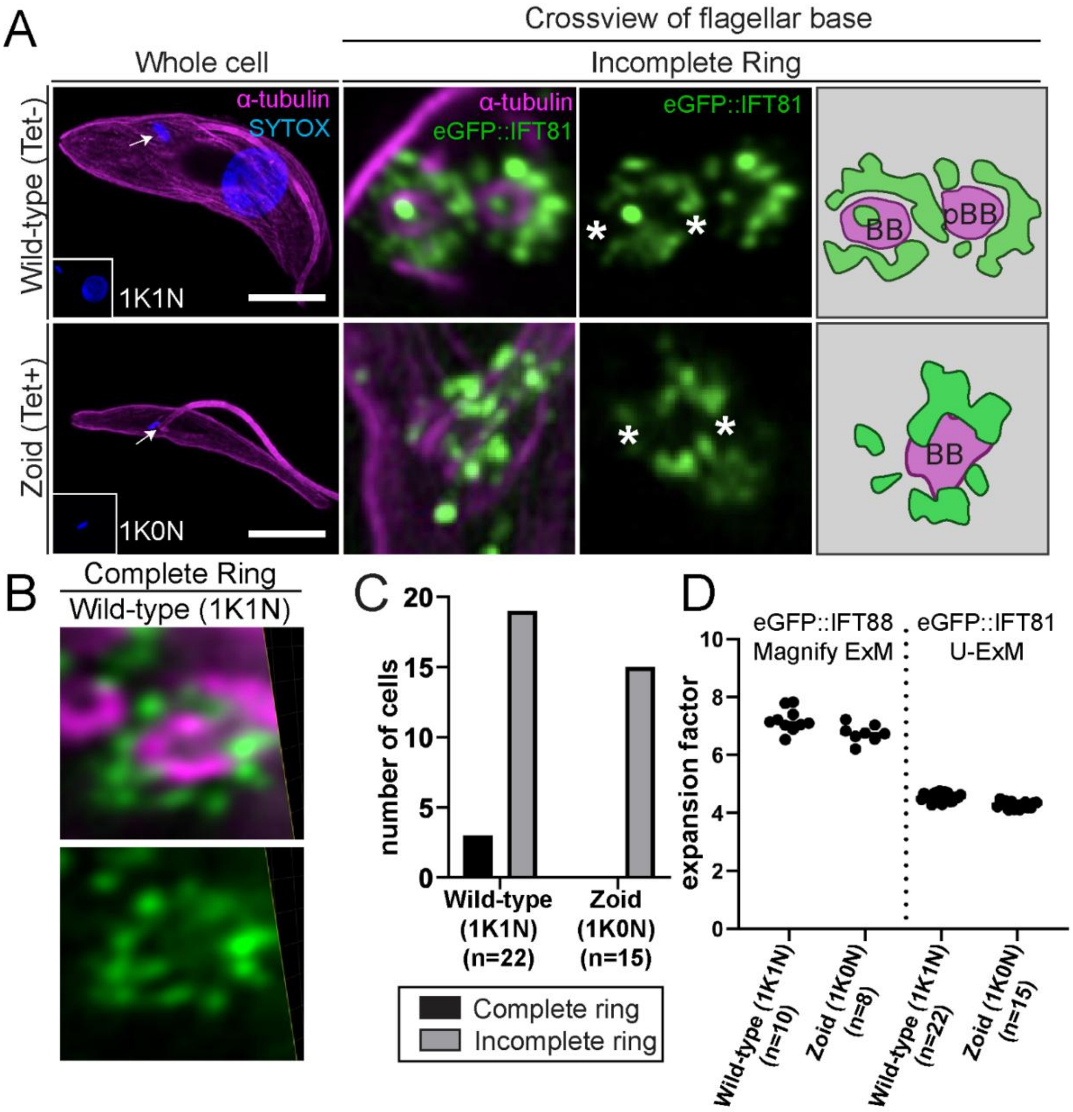


**Supplemental Figure 6: Ultrastructural Expansion Microscopy (U-ExM) reveals incomplete IFT occupancy patterns of eGFP::IFT81 at the flagellar base. (A)** U-ExM imaging of eGFP::IFT81 in non-dividing wild-type cells (1K1N) and zoids (1K0N). Scale bar 10 µm. Cross-views of the flagellum base (white arrows) show incomplete rings of IFT81 (white asterisks). Cartoon depiction (right) illustrates the distribution of IFT81 surrounding the basal body (BB) and the pro-basal body (pBB) when present. **(B)** Representative U-ExM image of eGFP::IFT81 in a wild-type cell showing a complete IFT occupancy ring. **(C)** Quantification of cells presenting complete or incomplete ring of IFT81 in non-dividing wild-type cells (1K1N) and zoids (1K0N). **(D)** Expansion factor measurements for Magnify ExM and U-ExM in wild-type cells and zoids. Expansion factors were calculated for each cell as the ratio of expanded to native axoneme diameter (220 nm). Magnify ExM achieved average expansion factors of 7.2x in wild-type cells and 6.7x in zoids, where U-ExM achieved 4.6x in wild-type cells and 4.3x in zoids.

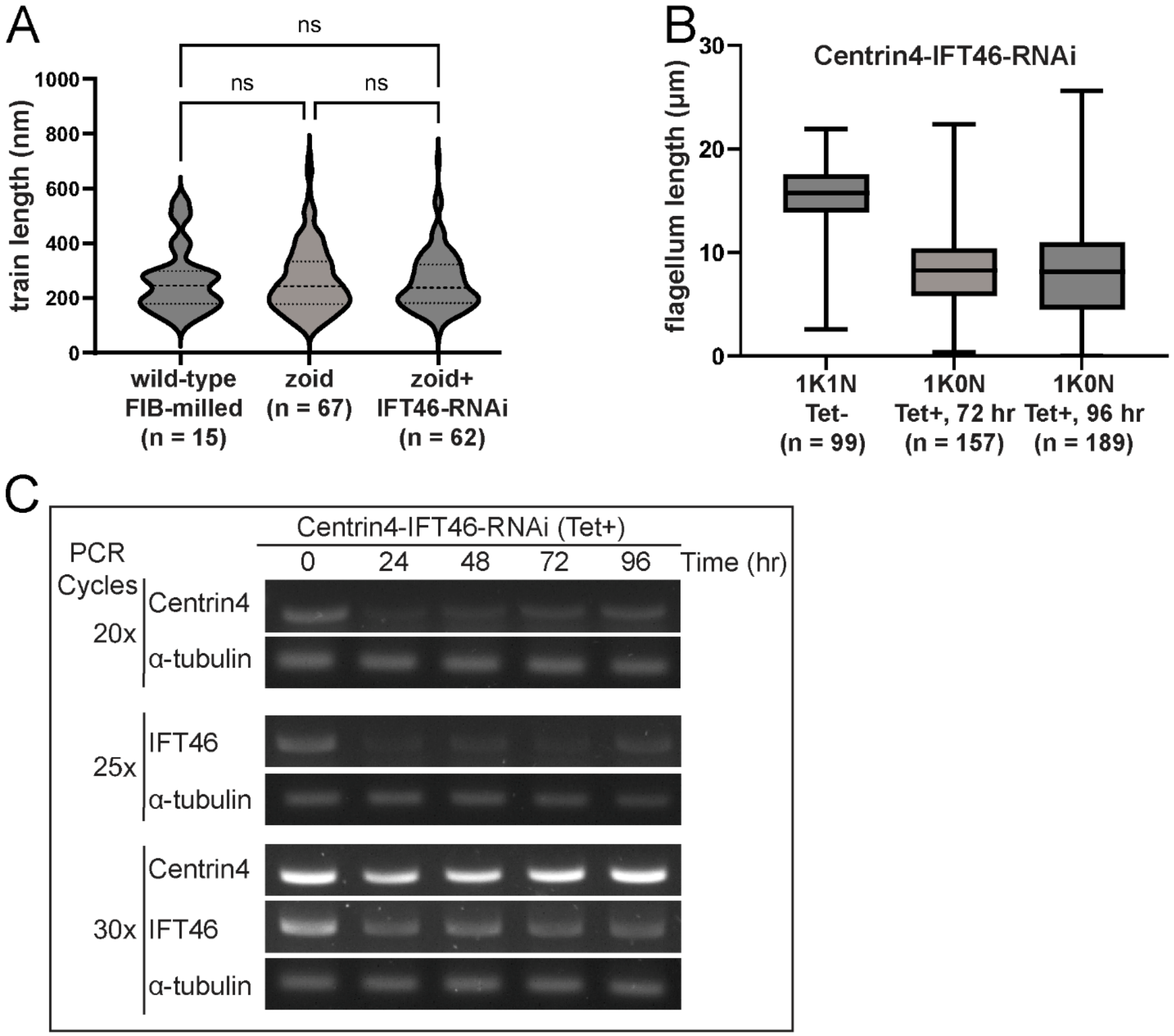


**Supplemental Figure 7: Characterization of IFT46 knockdown and its effects on IFT train and flagellum length. (A)** Lengths of initial-landing IFT trains measured from 2D tomogram slices were comparable across conditions, with no significant differences between FIB-milled wild-type cells (265 ± 118 nm), zoid (268 ± 123 nm), and IFT46-depleted zoids (261 ± 108 nm) (not significant (ns), p-value > 0.05). **(B)** Flagellum length measured by PFR immunofluorescence staining decreased upon dual knockdown of Centrin4 and IFT46. Average lengths were 15.6 ± 3.02 μm in uninduced, non-dividing (1K1N) cells, 8.27 ± 4.07 μm in zoids at 72 hours post-induction, and 7.84 ± 4.58 μm in zoids at 96 hours post-induction. **(C)** RT-qPCR analysis confirmed efficient depletion of both Centrin4 and IFT46 after 96 hours of tetracycline-induced RNAi.

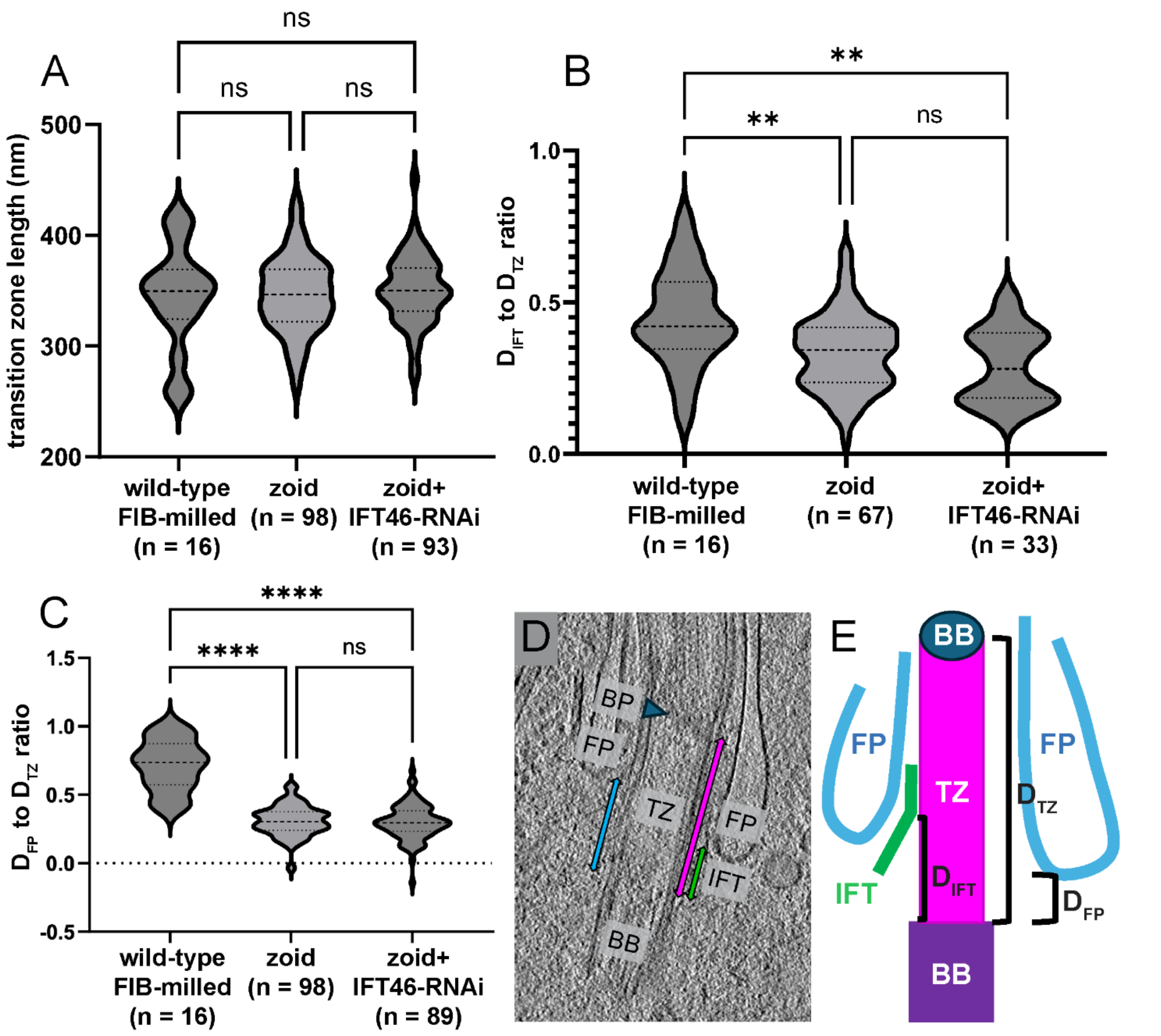


**Supplemental Figure 8: Altered flagellar pocket (FP) positioning in zoids correlates with proximal IFT docking along the transition zone. (A)** The length of the transition zone measured from 2D tomogram slices was comparable in FIB-milled wild-type cells (345 ± 46.4 nm), zoid (347 ± 34.2 nm), and IFT46-depleted zoids (351 ± 31.3 nm) . **(B)** The relative position of initial landing IFT trains, defined by the ratio of the distance from the landing site ($D_{IFT}$) to the proximal basal body over the length of the transition zone ($D_{TZ}$), was lowered in zoids (0.34 ± 0.12) and IFT46-depleted zoids (0.29 ± 0.12) compared to wild-type (0.45 ± 0.16). **(C)** The relative position of the FP membrane along the transition zone, defined by the ratio between the distance from the bottom of the FP to the proximal of the basal body ($D_{FP}$) over the full length of the transition zone ($D_{TZ}$), was lowered in zoids (0.31 ± 0.11) and IFT46-depleted zoids (0.29 ± 0.15) compared to wild-type (0.71 ± 0.19). **(D)** Representative tomogram slice of a FIB-milled wild-type cell showing measurement parameters. **(E)** Schematic diagram summarizing definitions of $D_{FP}$, $D_{IFT}$ and $D_{TZ}$.

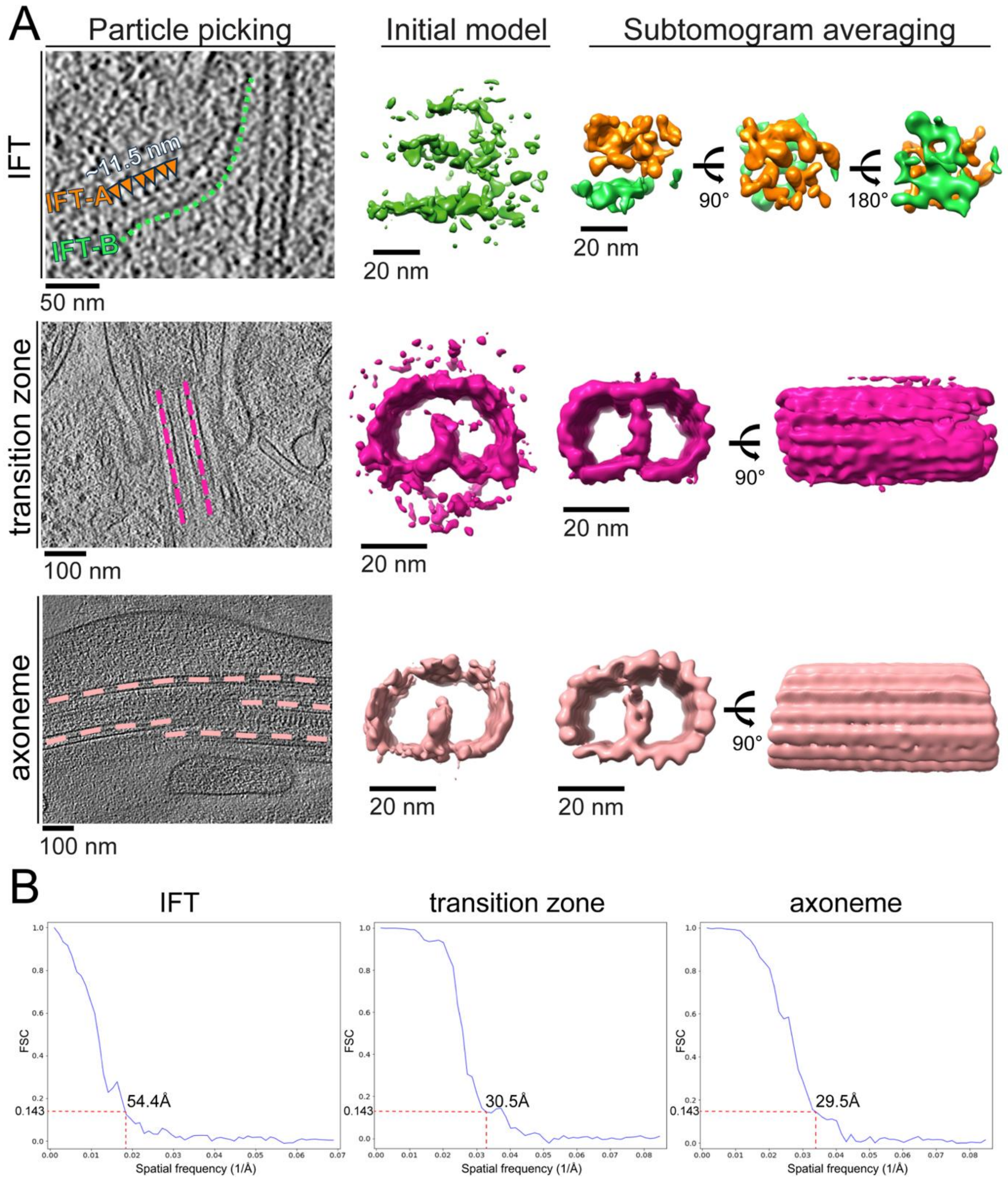


**Supplemental Figure 9: Summary of subtomogram averaging workflows and resolutions. (A)** Overview of particle processing for three structural groups: IFT train segments from zoids (top row), transition zone MTD (middle row), and axoneme MTD from FIB-milled wild-type cells (bottom row). For each group, representative particle extracted for *de novo* initial models, and final subtomogram averages are shown. **(B)** Masked Fourier shell correlation (FSC) curves for each averaged structure, with resolutions determined at the 0.143 FSC criterion.

**Movie S1:** Tomographic reconstruction and 3D segmentation of cryo-FIB-milled wild-type cells enabled the visualization of IFT train at the flagellar base. Scale bar 100 nm.

**Movie S2:** A Magnify-expanded wild-type cell revealed the incomplete occupancy of eGFP::IFT88 at the flagellar base.

**Movie S3:** A Magnify-expanded zoid revealed the incomplete occupancy of eGFP::IFT88 at the flagellar base.

**Movie S4:** Tomographic reconstruction and 3D segmentation of zoid flagellar base enabled the visualization of IFT train at the flagellar base. Scale bar 100 nm.

**Movie S5:** Comparative averaged cryo-EM maps of bidirectional IFT trains in *Chlamydomonas* and anterograde trains in *T. brucei*.

**Movie S6:** Tomographic reconstruction and 3D segmentation of a IFT46-depleted zoid revealed the wider IFT trains at the flagellar base. Scale bar 100 nm.
